# Thermo-Responsive Polymers Targeting Inflammation in Murine Colitis

**DOI:** 10.1101/2023.12.28.573545

**Authors:** Sufeng Zhang, Amy T. Jin, Wen Tang, Rachel Y. Zhang, Lihong Jing, Yixuan Zhou, Heng Zhang, Jochen K. Lennerz, Joshua R. Korzenik, Robert Langer, Giovanni Traverso

## Abstract

Targeting the site of inflammation is an ideal approach for treating inflammatory bowel disease (IBD). Inflammation targeting enables maximal drug-on-target effects while minimizing off-target side effects. Negatively charged drug carriers have been shown to facilitate drug delivery to inflamed colon mucosa after local administration. To modulate the negative charges and integrate responsiveness to stimuli, here we describe thermo-responsive, inflammation-targeting (TRIT) hydrogels based on functionalized poly(*N*-isopropylacrylamide-*co*-methacrylic acid) (PNIPAM-MAA). We show that both chemical modification types and polymer molecular weights affect the resultant microgels’ adhesion to the inflamed colon in dextran sulfate sodium (DSS)-induced murine colitis *in vivo*. Further, we quantified the correlations between microgels’ adhesion and colitis severity for individual mice, demonstrating that the microgels’ adhesion correlated directly with weight loss percentage in DSS-treated mice. By exploiting charge-mediated interaction and thermo-responsiveness, TRIT hydrogels represent a promising strategy to target inflamed colon mucosa as a drug delivery platform for colonic IBD treatment.

**Teaser:** This study developed thermo-responsive, inflammation-targeting (TRIT) hydrogels that harness charge-mediated interaction and sol-to-gel transition to target inflamed colon mucosa as a new approach for treating inflammatory bowel disease.

## Introduction

Inflammatory bowel disease (IBD) in its two major clinically defined forms, Crohn’s disease (CD) and ulcerative colitis (UC), is a chronic, idiopathic inflammatory set of conditions that may affect the entire gastrointestinal (GI) tract (*1*). The etiology of IBD is unclear, though it is commonly understood that certain environmental factors trigger an inappropriate inflammatory response to intestinal microbes in genetically susceptible hosts (*2*). Recently, IBD has evolved into a global disease with rising prevalence in many countries (*3*). There is no cure for IBD, necessitating life-long medication for patients. Frequent and long-term uses of systemic immunosuppressive drugs are associated with side effects and serious complications, such as opportunistic infections, malignancies, and liver toxicity (*4*). Drug delivery methods that selectively target the inflamed intestine would increase drug efficacy and minimize the exposure of healthy or distant tissues to immunosuppressive drugs, reducing side effects.

Under inflammation, the colonic mucus layer that normally covers the epithelium tends to decrease in thickness or become depleted at the inflamed region (*5, 6*). The inflamed microenvironment at the damaged mucosa contains a broad spectrum of pathophysiological features that could be exploited for the rational design of targeted drug delivery, using size-, charge-, ligand-receptor-, degradation-, or microbiome-mediated strategies (*7*). The charge-mediated targeting strategy relies on the accumulation of positively charged proteins at the inflamed colonic mucosa, which provide an instructive cue for negatively charged drug carriers to interact with. Previously, we developed a negatively charged, small molecule-based hydrogel that preferentially targeted the inflamed colon for localized dexamethasone delivery in IBD (*8*). This delivery technique improved drug efficacy and reduced systemic drug exposure in rodent models of IBD.

To enhance drug delivery targeting the inflamed colon, we expanded the small-molecule-based hydrogel to polymeric hydrogels with *in-situ* gelation using thermo-responsive materials. Thermo-responsive polymers have been widely used for biomedical applications for injection convenience as a liquid and subsequent gelation at the injection site to provide local drug delivery and release (*9–11*). Poly(*N*-isopropylacrylamide) (PNIPAM)-based materials are synthetic thermo-responsive polymers that have been extensively studied, due to their temperature-sensitive phase change, inert stability, and good biocompatibility (*12–15*). These polymers undergo a sharp coil-globule transition and phase separation at their lower critical solution temperature (LCST) in water, during which the interaction of hydrophobic components in the polymers forms nanodomains.

PNIPAM-based thermo-responsive polymers have been used as scaffolds for cell culture models of human intestinal epithelium (*16*) and drug delivery systems for growth factors in wound healing and small-molecule therapeutics for cancer therapies or photothermal therapies (*17–19*). Their high-water content and lack of chemical crosslinks provide good biocompatibility and facilitate clearance from the body. Administration via enemas in the liquid format of these thermo-responsive polymers allows broad dispersion in the colon, followed by *in-situ* gelation to localize therapeutics to ulcers for maximal efficacy. A thermo-sensitive delivery platform was reported using copolymers consisting of polyethylene glycol and polypropylene glycol for budesonide and mesalamine delivery in rodent IBD models (*20*). The delivery platform improved drug efficacy and showed greater colonic retention of drugs than drug liquids; however, detailed characterization of the material’s physical properties was lacking. While using thermo-responsive materials for drug delivery to colonic IBD is advantageous and promising, there are few studies on such delivery systems for this application.

Here, we report the development and characterization of PNIPAM-based thermo-responsive polymers targeting the inflamed colonic mucosa in murine models of colitis. We hypothesize that negatively charged PNIPAM-based polymers target the site of inflammation via charge-mediated interaction and subsequently form drug depots at the inflamed colonic mucosa through the sol-to-gel transition. Taking advantage of their physicochemical characteristics suitable for local drug delivery in colonic IBD, these thermo-responsive, inflammation-targeting (TRIT) polymers could have significant applications in IBD treatment. In this work, we choose PNIPAM-MAA for chemical modification because it responds sharply to temperature change (*21*) and possesses carboxylic acids for reaction. To tune the charges of PNIPAM-MAA, we conjugated two different molecules to PNIPAM-MAA, respectively: taurine (S modification) to increase the negative charges and *N,N*-dimethyl-ethylenediamine (N modification) to decrease the negative charges. The sulfonates in the S modification are strongly hydrated anions, while the tertiary amines in the N modification are weakly hydrated cations. The negative charges by sulfonates from the conjugated taurine are considered stable under acidic pH associated with intestinal inflammation (*22*), compared to the pH-dependent carboxylic acids. Furthermore, taurine is known to have antioxidant, antimicrobial, and anti-inflammatory effects that could benefit IBD treatment (*23–27*).

PNIPAM-MAA of two molecular weights, 10 kDa and 60 kDa, were employed for the chemical modifications. Using PNIPAM-MAA as a control, the modified polymers’ size and zeta potential were characterized before and after the gelation *in vitro*. We then evaluated the adhesion of these polymers to the inflamed colon using dextran sulfate sodium (DSS)-induced acute colitis in mice *in vivo*, compared to healthy mice. The selection of DSS-induced acute colitis to study polymers targeting the inflamed colon was due to the quick onset of inflammation and the well-established experimental procedure in generating this colitis model (*28, 29*). Owing to the heterogeneous nature of chemically induced colitis, we further analyzed correlations between polymer targeting the inflamed colon and colitis parameters, including body weight loss percentage, colon length, colonic myeloperoxidase (MPO) activity, and histology scores from individual mice. Examining polymers’ mucosal binding with colitis parameters could provide a further understanding of interactions between polymeric hydrogels and the biological microenvironment in colitis. These analyses may also advance our knowledge of comparing materials’ binding capacity with chemically induced colitis models.

## Results

### Functionalization of PNIPAM-MAA with the S and N ligands

We chose PNIPAM-MAA for chemical modification, owing to the availability of carboxylic acids for reaction and the sharp response of PNIPAM-MAA to temperature change to undergo phase transition (*21*). The molar content of MAA in the PNIPAM-MAA we used was 10%. This was chosen because higher percentages of acrylic acids cause the cloud point to disappear due to sufficient solubility of MAA that offsets the aggregation of the hydrophobic temperature-sensitive components (*30*). We employed two different ligands to functionalize PNIPAM-MAA by EDC/NHS chemistry; one is taurine (S modification) to increase the negative charge, and the other is *N,N*-dimethyl-ethylenediamine (N modification) to reduce the negative charge (Fig. 1A). These reactions were performed for PNIPAM-MAA of two molecular weights (*M*_n_), 10 and 60 kDa. The resultant polymers were termed 10KS, 60KS, 10KN, and 60KN, with the unmodified polymers 10K and 60K as controls. We chose taurine to modify PNIPAM-MAA because taurine is a naturally occurring amino sulfonic acid, and the ionization of sulfonic acids is stable compared to the pH-dependent ionization of carboxylic acids. This S modification enabled higher negative charges on 10KS and 60KS than the corresponding 10K and 60K in the inflamed microenvironment, where inflammation likely increases the local acidity in the niche. During the reaction, we used excessive EDC to activate PNIPAM-MAA and excessive S or N ligands for maximal substitution on PNIPAM-MAA; the feed molar ratio was 1:3:3:10 between MAA, EDC, NHS, and S or N ligand for modification. The results showed a substitution percentage of 33 - 40% for S modification (40.5% for 10KS and 33.3% for 60KS) and about 19% for N modification (19.1% for 10KN and 19.2% for 60KN) on PNIPAM-MAA (Fig. 1B), calculated based on the ^1^H-NMR spectra (fig. S1).

**Fig. 1.**
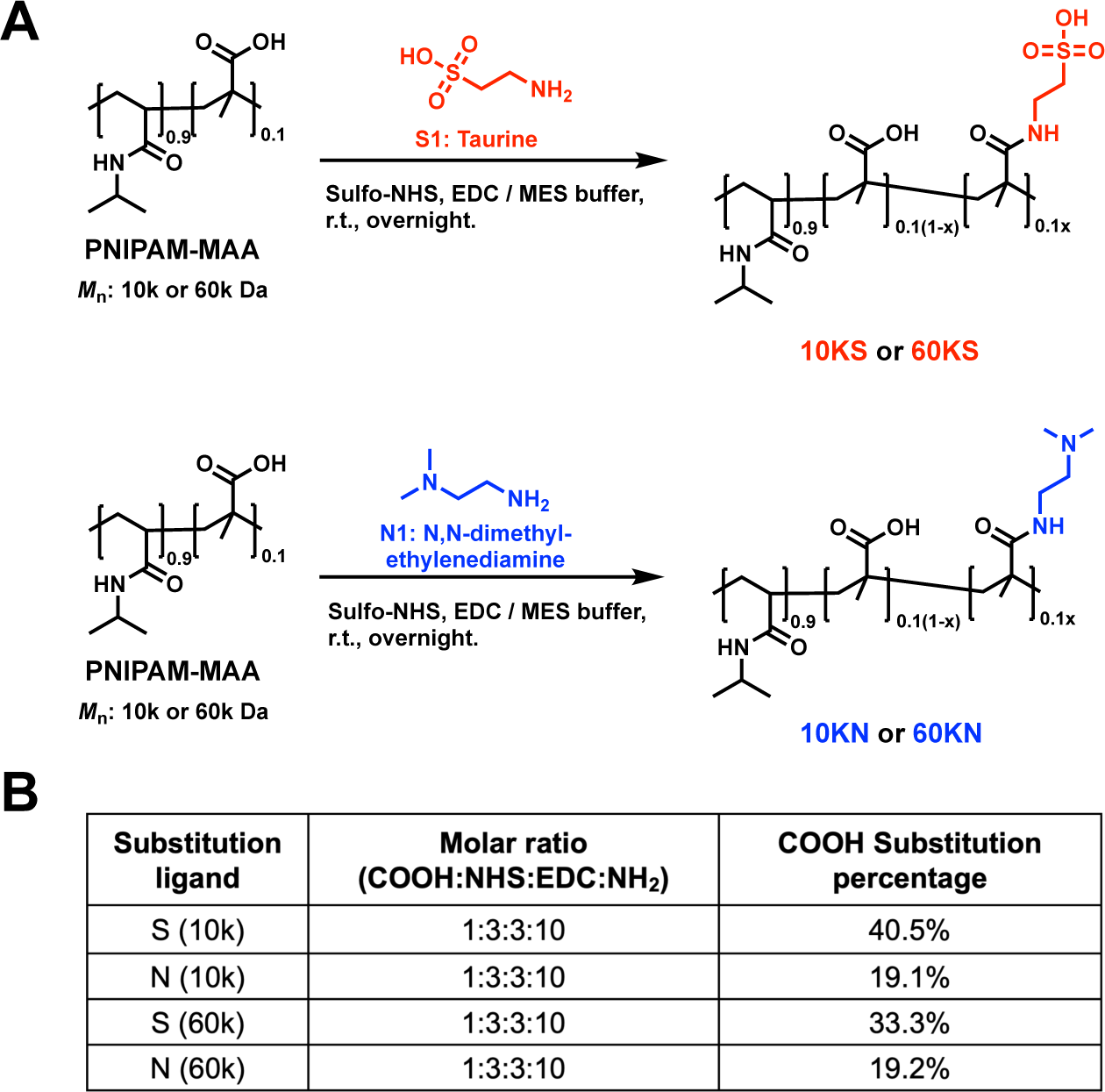
Chemical modifications of PNIPAM-MAA. PNIPAM-MAA (10% MAA molar content) of 10,000 (10K) and 60,000 (60K) g/mol in molecular weight were used for the chemical modification. The carboxylic acids on PNIPAM-MAA were functionalized with taurine (S modification) or *N, N*-dimethyl-ethylenediamine (N modification). **(A)** Synthesis of the S or N modified PNIPAM-MAA using the EDC/NHS chemistry. After synthesis, the resulting mixture was dialyzed against 0.2 M carbonate buffer (x6), water (x3), and then lyophilized. The molecular weight cutoff (MWCO) of the dialysis tubing was 3.5 kDa for the 60K polymers and 1.0 kDa for the 10K polymers. **(B)** The structure of the modified polymers was confirmed by ^1^H-NMR, and the substitution percentage of S or N modification on PNIPAM-MAA (10K) and PNIPAM-MAA (60K) was determined, respectively.

### Cloud Point temperature and microgels’ morphology of the functionalized PNIPAM-MAA

First, we characterized and compared the LCST behavior of PNIPAM-MAA and the modified polymers using the cloud point temperature (*T*_cp_), measured by turbidimetry (*31*). We determined the *T*_cp_ of 10K, 10KS, 10KN, 60K, 60KS, and 60KN of various concentrations. The Transmittance% (T%) curves of all polymers at 5, 10, 15, 25, and 35 mg/ml were measured as a function of temperature ranging from 25°C to 50°C (fig. S2A). The *T*_cp_ of each polymer was calculated by determining the inflection point of the T% curve and plotted as a function of polymer concentration (fig. S2B). Further, the *T*_cp_ of each polymer at different concentrations was compared (fig. S2C), and the correlation between *T*_cp_ and polymer concentration for PNIPAM-MAA and modified polymers were fitted using a four-parameter fit curve (Fig. 2A). The results showed that both S and N modifications elevated the *T*_cp_, compared to the PNIPAM-MAA, with the S modification being more pronounced for both 10K and 60K PNIPAM-MAA. Since the S modification introduced sulfonic acids into the polymer, which increased the hydrophilicity of the polymer, it is expected that the S modification elevated the *T*_cp_. On the other hand, replacing acrylic acids with N also increased *T*_cp_, which could be attributed to the increase in hydrophilicity due to the addition of tertiary amine into the polymer. From the four-parameter fitted curves of *T*_cp_ vs. polymer concentration, the critical gelation concentration (CGC) of each polymer at *T*_cp_ of the human body temperature 37°C was determined, which were 5.6, 38.7, and 32.4 mg/ml for 10K, 10KS, and 10KN, and 4.9, 16.5, and 10.1 mg/ml for 60K, 60KS, and 60KN, respectively. Each polymer was then prepared at its corresponding concentration, T% curves were measured, and experimentally confirmed individual’s *T*_cp_ within 37 ± 0.5°C (Fig. 2B, 2C).

**Fig. 2.**
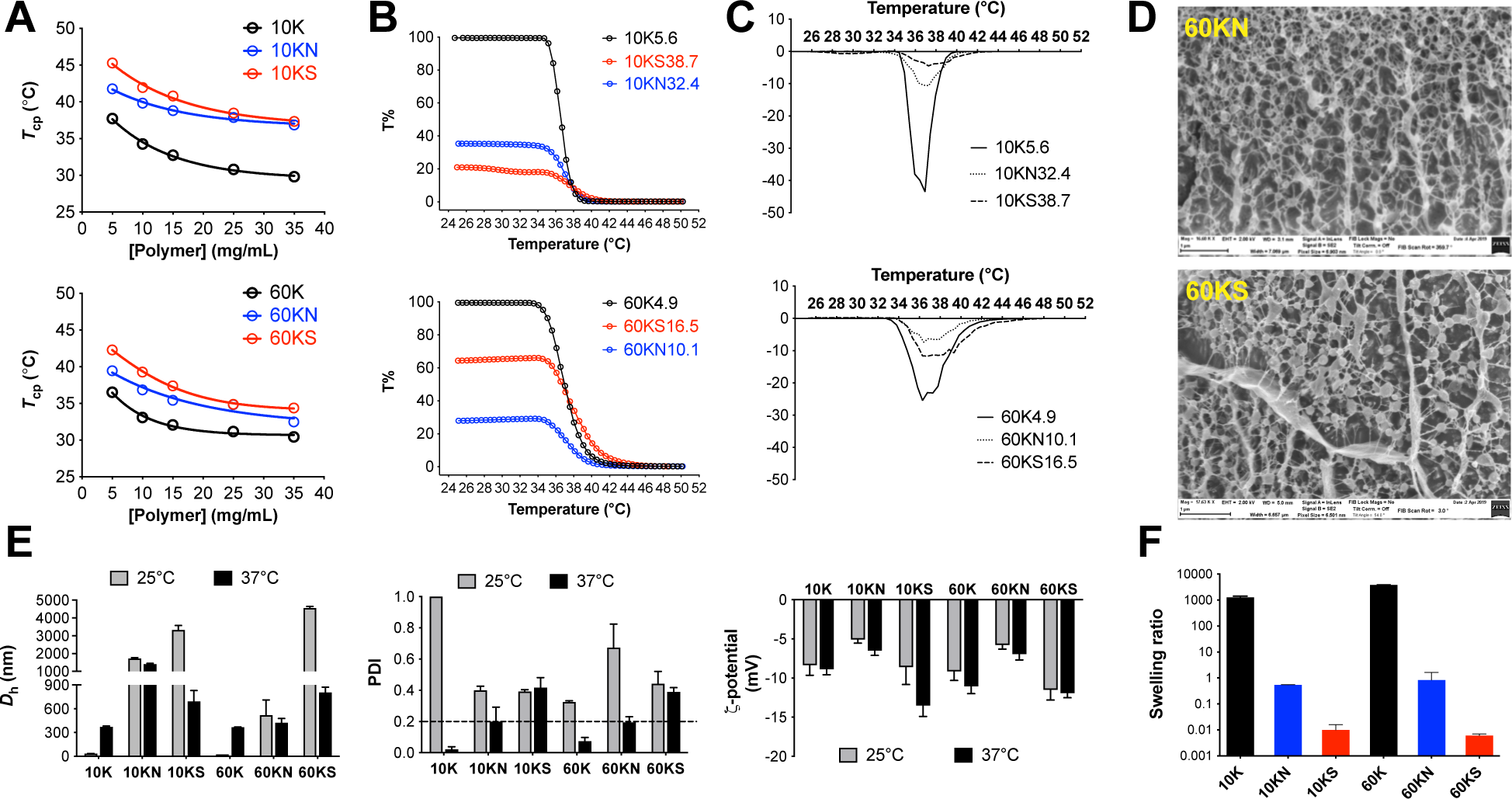
Characterization of chemically modified PNIPAM-MAA (10KN, 10KS, 60KN, and 60KS) using the unmodified polymers (10K and 60K) as control. **(A)** The lower critical solution temperature (LCST) of all polymers (10K, 10KN, 10KS, 60K, 60KN, and 60KS) was determined using the cloud point method (*T*_cp_) and measured at different polymer concentrations (circles indicate experimental data). A four-parameter fit curve was plotted to correlate the polymer concentration with *T*_cp_ (lines indicate the simulated curves calculated based on the four-parameter curve fit). (**B**) Transmission percentage (T%) curves of polymers at concentrations of *T*_cp_ = 37°C. For 10K, 10KN, and 10KS, the concentration of *T*_cp_ = 37°C for each polymer is 5.6 mg/ml, 38.7 mg/ml, and 32.4 mg/ml; For 60K, 60KN, and 60KS, the concentration of *T*_cp_ = 37°C for each polymer is 4.9 mg/ml, 16.5 mg/ml and 10.1 mg/ml. (**C**) First derivatives of the T% curves as a function of temperature for 10K, 10KN, 10KS and 60K, 60KN, 60KS at the concentration of *T*_cp_ = 37°C for each polymer. The maximal point of the first derivative curve of each line indicated the *T*_cp_. The results confirmed experimentally that the polymer concentration from the four-parameter fit curve led to *T*_cp_ = 37°C for each polymer. (**D**) Representative cryo-scanning electron microscopic (Cryo-SEM) images of 60KN and 60KS. To ensure gelation, polymers were used at 2x the concentration of *T*_cp_ = 37°C. **(E)** The hydrodynamic size (*D*_h_) using Z-average intensity mean, polydispersity index (PDI), and zeta potential (ζ-potential) measurements of all polymers at their concentrations of *T*_cp_ = 37°C under 25°C and 37°C, respectively. Data are Mean ± SD. **(F)** The swelling ratio for each polymer was determined by the ratio of spheric volumes of each polymer calculated from their size measurement under 37°C and 25°C. Data are Mean ± SEM.

The internal structure of the 60KS and 60KN polymeric hydrogels at the micro/nanoscale was examined and visualized by cryo-scanning electron microscopy (cryo-SEM). Representative images showed the morphology of the spherical particles in the microgels (Fig. 2D). As a traditional approach, snap-freezing may have limitations in the cryo-SEM imaging process. The “wall-like” structures in these images were probably artifacts of ice crystals during the sublimation process in the sample preparation (*32*). However, these images roughly indicated the internal structure of these modified thermo-responsive polymers after gelation at the micro/nanoscale, which represented an initial step in understanding the *in-situ* structure of these polymers when applied *in vivo*.

### Hydrodynamic size and zeta potential of the functionalized PNIPAM-MAA

Next, we measured the hydrodynamic diameter (*D*_h_) and zeta potential (ζ-potential) of PNIPAM-MAA and the modified polymers. *D*_h_ reveals the swelling or shrinking behavior of these polymers in response to temperature change, while the ζ-potential is critical in confirming the S and N modifications and understanding the effect of surface charge on polymers’ binding to the inflamed colon. First, we measured *D*_h_, polydispersity index (PDI), and ζ-potential of the six polymers, 10K, 10KN, 10KS, 60K, 60KN, and 60KS at 0.5 mg/ml in PBS (1×, pH = 7.4) as a function of temperature (fig. S3). Both 10K and 60K showed a sharp increase in *D*_h_ when the temperature was raised to 32 −34°C and reached a plateau after 37°C. In contrast, the modified polymers showed relatively similar *D*_h_ across the temperature range. A gradually decreased PDI was observed for all polymers upon increasing temperature. There was an increase in the negative ζ-potential for 10K and 60K as the temperature increased. Importantly, when the temperature was above 37°C, 10KN and 60KN showed the least negative ζ-potential, followed by 10K and 60K. 10KS and 60KS remained the highest negative ζ-potential across the temperature range.

To understand the polymer’s behavior under physiological temperature, we characterized *D*_h_ and PDI of the polymers as a function of temperature at each polymer’s CGC of *T*_cp_ = 37°C (fig. S4). For 10K, 10KN, and 10KS, the concentration of each polymer at *T*_cp_ = 37°C is 5.6, 32.4, and 38.7 mg/ml in PBS (1×, pH = 7.4). For 60K, 60KN, and 60KS, the concentration of each polymer at *T*_cp_ = 37°C is 4.9, 10.1, and 16.5 mg/ml in PBS (1×, pH = 7.4). Upon temperature increased to 32°C, 10K and 60K started to show an increase in *D*_h_ while the modified polymers displayed a reduction in *D*_h_. The decrease in PDI was more advanced for 10K and 60K than the modified polymers. However, the ζ-potential of individual polymers at the CGC of *T*_cp_ = 37°C could not be obtained due to the blackened gold-plated copper electrodes during the measurement, which was associated with the high polymer concentrations and the extended measurement duration required for the temperature range. Therefore, we measured and compared individual polymers’ *D*_h_, PDI, and ζ-potential at their CGC of *T*_cp_ = 37°C under 25°C and 37°C, respectively (Fig. 2E). The results showed that both 10K and 60K formed microgel particles under 37°C with a *D*_h_ of 373.2 nm and 367.7 nm (PDI < 0.2), compared to 34.4 nm and 23.5 nm at 25°C (PDI >> 0.2). At 37°C, the N modification led to a smaller PDI than the S modification, while the S modification showed a more considerable decrease in *D*_h_ than the N modification, compared to that at 25°C. Although shrinking from 25°C to 37°C, the *D*_h_ of the modified polymers was still larger than that of PNIPAM-MAA, being 692.6 nm for 10KS, 1414.3 nm for 10KN, 373.2 nm for 10K, and 806.1 nm for 60KS, 425.2 nm for 60KN, 367.7 nm for 60K. When comparing ζ-potential, 10KN and 60KN showed the least negative charge, 10K and 60K in the middle, and 10KS and 60KS the highest negative charge of all polymers at both 25°C and 37°C, which also confirmed the successful conjugation of ligands onto PNIPAM-MAA.

When the temperature was raised from 25°C to 37°C, 10KN or 60KN demonstrated a ∼1.2-fold decrease in *D*_h_, while 10KS and 60KS both showed a ∼5.2-fold decrease in *D*_h_. Assuming spherical shapes of the formed particles and using the measured *D*_h_, individual polymers’ swelling/shrinking behaviors were calculated as the volume ratio of the particles measured at 37°C and 25°C (Fig. 2F). The result showed a highly swelling behavior of about 1000 times after gelation for 10K and 60K, while 10KS and 60KS demonstrated the highest shrinking behavior of about 100 times. 10KN and 60KN were in the middle, shrinking roughly 0.5 times. Presumably, 10K and 60K possessed more carboxylic acids than the modified polymers, which were available to form hydrogen bonding and absorb water during the gelation, thereby leading to the swelling of the PNIPAM-MAA gel. On the other hand, the S and N modifications reduced the number of carboxylic acids for hydrogen bonding and increased the influence of hydrophobic collapse, leading to a compact structure. The high concentrations of the modified polymers of *T*_cp_ = 37°C might also contribute to particle shrinkage due to inter-microgel steric compression and ion-induced deswelling (*33, 34*).

### Adhesion of PNIPAM-MAA and modified polymers to the inflamed colon in mice *in vivo*

The *in vivo* studies were conducted with two main goals: (1) To assess the selective targeting of polymers to the inflamed colon, compared to the healthy colon *in vivo*; (2) To determine the effect of negative charge on polymer’s targeting the inflamed colon by comparing the S modification with the N modification and PNIPAM-MAA. Texas Red^®^ dextran (Ex = 570 nm, Em = 620 nm) was mixed with polymers for *in vivo* tracking. Considering that a small-molecule dye may escape from the hydrogel, we chose Texas Red^®^ dextran of 10 kDa in molecular weight to mix with the polymers for rectal administration. We chose a low concentration of the dye, 0.1 mg/ml, to mix with individual polymers to minimize any effect of the dye on the polymers’ gelation. The same amount of the dye was used for all individual polymers. The molar ratio of the polymers to the dye ranged from 8.2 to 387. The mixture of each polymer with the dye was characterized by size and ζ-potential measurement at 25°C and 37°C, respectively (fig. S5). The result showed a similar pattern in the size and ζ-potential measurement for the mixture of the dye with individual polymers as observed in the polymers alone, indicating that the addition of the dye did not significantly alter the gelation. However, we noticed that the dye alone was negatively charged and formed nanoparticles of about 200 nm with a PDI of 0.23.

Next, we determined the selective targeting of polymers to the inflamed colon by comparing the polymer retention in the inflamed colon with the healthy colon. All polymers, 10K, 10KN, 10KS, 60K, 60KN, and 60KS, and the dye alone, were evaluated. Mice were given 3% DSS in drinking water for five consecutive days and then switched to regular drinking water for another two days. On day 7, the mixture of each polymer with the dye was administered rectally to colitic mice and healthy mice, respectively. Colons were dissected at 6h post-enema, measured in length, imaged by IVIS, and analyzed by histology and MPO assays (Fig. 3A). The DSS-induced acute colitis in mice was monitored by the body weight loss percentage (weight loss%) and the colon length (fig. S6). All DSS-treated mice showed a significant reduction in body weight and shorter colon length than the healthy controls. For all polymers, each group contained six mice except the healthy controls for 60KN, in which there were three mice because the other three mice were accidentally sacrificed at 3h post-enema. The 3 cm distal colon from individual mice treated with different polymers and the dye-alone were imaged by IVIS (Fig. 3B); full IVIS images with detailed descriptions were also shown in fig. S7. The quantification of IVIS fluorescence showed that all polymers retained significantly higher fluorescence in the colitic colon than in the healthy colon, including the dye alone (Fig. 3C). After IVIS imaging of the 3 cm distal colon (denoted “Distal”), 1 cm colon tissue was used for MPO analysis, 1 cm for histology assessment, and 1 cm for homogenization to quantify the fluorescence using a microplate reader. Moreover, 2 cm colon tissue above the imaged distal colon (denoted “Proximal”) was used for additional MPO and histology assessment. A significant increase in MPO activity was observed in the inflamed colon compared to the healthy colon groups (figs. S8, S9). The DSS-treated groups also showed significantly higher histology scores than the healthy controls (figs. S10, S11), and the histology images of individual colon tissues are shown in fig. S12.

**Fig. 3.**
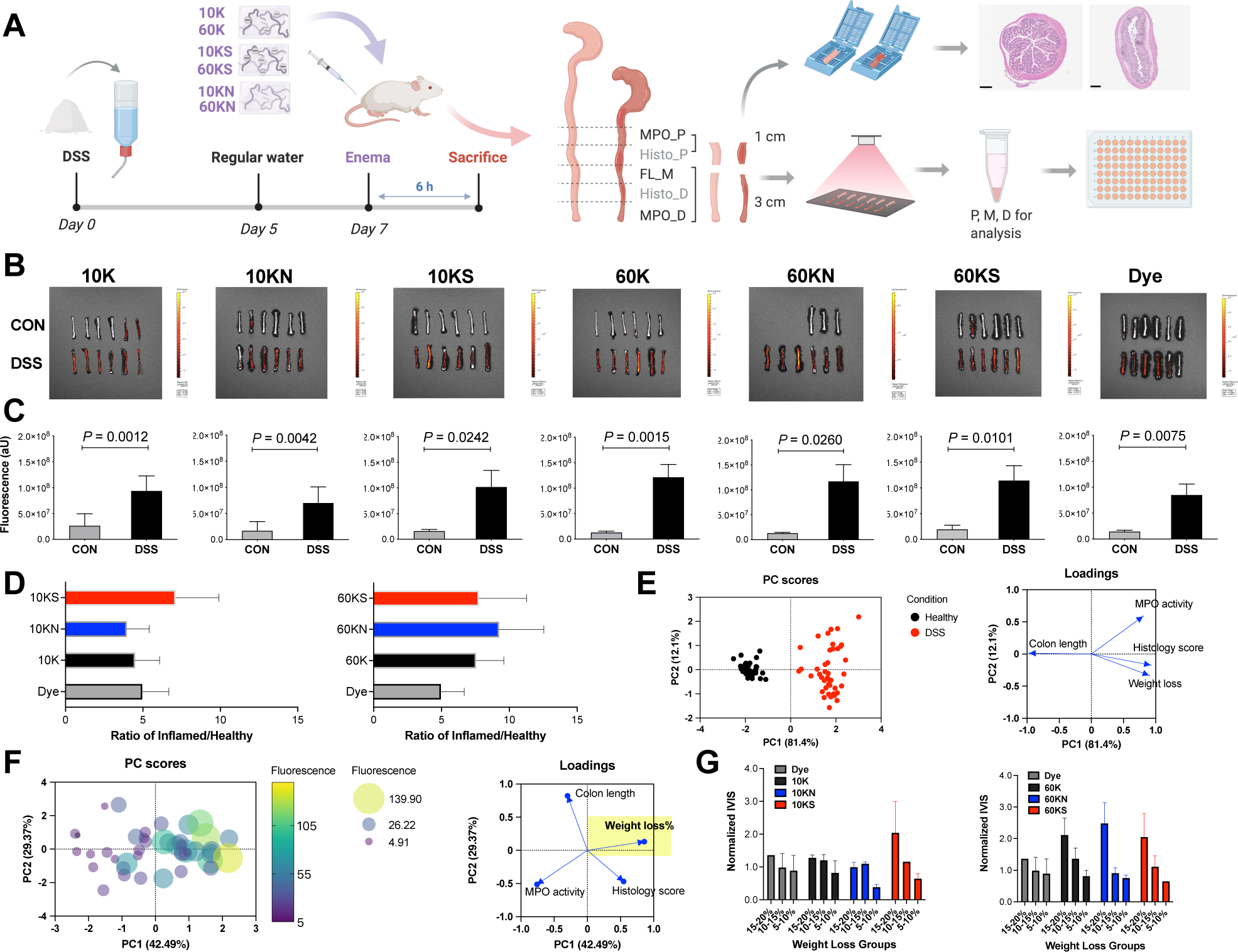
*In vivo* evaluation of polymers’ adhesion to the inflamed colon and their correlations with colitis parameters. A fluorescent dye, Texas Red^®^ dextran, was mixed with each polymer for *in vivo* tracking. The concentration of the dye used was 0.1 mg/ml for all polymers. **(A)** A schematic outlining the *in vivo* evaluation of polymers. Representative histology images showed the colon tissue from colitic mice and healthy controls. P: proximal, M: middle, D: distal. **(B)** IVIS images of the colon tissue from DSS-treated mice and healthy controls for the evaluation of all polymers and the dye alone. CON: healthy controls; DSS: DSS-treated mice. **(C)** Fluorescence quantification of IVIS images in **(B)**. **(D)** The ratios of fluorescence intensity in the colon tissue of the DSS-treated and healthy mice (Ratio of Inflamed/Healthy) were calculated for each polymer and the dye alone. **(E)** Principal component analysis (PCA) was performed on all mice with the loadings of weight loss percentage (%), histology score, colon length, and MPO activity. Healthy and DSS mice are clearly separated along the PC1 axis, which explains 81.4% of the variation. **(F)** PCA analysis of the spread of disease heterogeneity and its effect on polymer adhesion in DSS-treated mice. The variable loadings are weight loss%, histology score, colon length, and MPO activity. **(G)** IVIS fluorescence intensity as a function of weight loss% of mice in three bins for each polymer and the dye alone. Normalized IVIS was calculated by the IVIS fluorescence in the polymer groups divided by the IVIS fluorescence in the dye alone group.

To compare the binding capability of polymers, we calculated the ratio of fluorescence intensity for the inflamed colon divided by the healthy colon, denoted Ratio of Inflamed/Healthy, for all polymers and the dye alone. The ratio of Inflamed/Healthy was 7.12 for 10KS, 4.50 for 10K, 3.96 for 10KN, 7.76 for 60KS, 7.55 for 60K, 9.25 for 60KN, and 4.97 for the dye alone (Fig. 3D). 10KS showed the highest ratio of Inflamed/Healthy, compared to 10K, 10KN, and the dye alone. This observation matched our hypothesis that increased negative charge increased polymer adhesion to the inflamed colon. However, 60KN showed a higher ratio of Inflamed/Healthy than 60KS and 60K, and these three polymers were much higher than the dye alone. Since the same amount of Texas Red^®^ dextran was used across all groups, the difference between each polymer group and the dye alone indicates the targeting capability of individual polymers. The result showed that increased negative charges on PNIPAM-MAA enhanced the polymer targeting the inflamed colon for the low MW of 10 kDa; however, at the high MW of 60 kDa, there was little effect of charges on the polymers’ targeting the inflamed colon.

### Correlation between polymer adhesion to the inflamed colon and colitis parameters

Due to individual differences and variations in the susceptibility to DSS-induced colitis, heterogeneity of colitis severity in DSS-treated mice is expected (*28, 35*). To better visualize the data and gain insights, principal component analysis (PCA) was performed on all mice with the loadings of colitis parameters, including weight loss percentage, colon length, MPO activity, and histology score. Healthy and DSS mice were clearly separated along the PC1 axis, which explains 81.4% of the variation (Fig. 3E). This is expected as these variables are known biomarkers for the development of acute colitis. There is more spread along PC2, which includes contribution primarily from MPO activity but also from weight loss and histology score.

To further investigate the spread of disease heterogeneity and its effect on polymer adhesion, we investigated the correlation between the fluorescence retention of all polymers studied and the colitis parameters in DSS-treated mice. We quantified the colonic fluorescence retention using two methods: immediate direct quantification of the 3 cm distal colon by IVIS imaging (fig. S13A) and fluorescence measurement normalized against the total protein of the homogenized 1 cm colon tissue from the 3 cm colon after IVIS imaging (fig. S13B). We monitored the weight loss% of individual mice in DSS-treated groups and healthy controls (fig. S13C). The logarithm of fluorescence intensity, quantified by IVIS and homogenized colon tissue, was proportional to the weight loss% of all mice studied, displaying a linear relationship by Pearson’s correlation (fig. S13D, E; *P* < 0.05). We then analyzed the correlation between the fluorescence intensity and weight loss% for colitic mice only. The correlation between the logarithm of fluorescence intensity and weight loss% was significant for all 60K-based polymers and most 10K-based polymers (fig. S13F, G; *P* < 0.05). However, neither correlation was significant for the dye alone group (*P* = 0.1327 for IVIS and *P* = 0.2087 for homogenized colon tissue). Next, PCA was performed on the DSS-treated mice with variable loadings of weight loss percentage, colon length, MPO activity, and histology score (Fig. 3F). A correlation between fluorescence and PC1 can be seen, and a least squares regression indicated a significantly nonzero contribution (*P* < 0.0001) of weight loss to fluorescence. The colon length, histology score, and MPO activity contribute to PC2 (29.4% of variation), but they do not have a significant effect on fluorescence. The fluorescence intensity quantified from all mice was collectively plotted against individual colitis parameters, including weight loss percentage, colon length, MPO activity, and histology score (fig. S14). A linear relationship was shown between the logarithm of the fluorescence and the weight loss% for both the IVIS quantification and the fluorescence quantification from homogenized colon tissue. After separating the mice into bins based on weight loss%, we found that the difference in adhesion between polymers was more apparent at higher weight loss% (Fig. 3G). Thus, to lower the variation and spread of mouse disease conditions, we sought to compare the adhesion of polymers in mice whose weight loss was more than 15%.

To differentiate the binding capacity between the S and N modified polymers, we reduced the dye concentration from 0.1 to 0.02 mg/ml to eliminate potential effects from the dye. We also included a 70 kDa Texas Red^®^ dextran dye to mix with the 60 kDa polymers. We characterized the size, PDI, and ζ-potential of the polymers mixing with the dye, and compared the mixtures when the dye was at 0, 0.02, and 0.1 mg/ml under 37°C (fig. S15). The results showed that mixing with a dye led to a more compact size and smaller PDI of the microgels, and a higher concentration of the dye enhanced such an effect. Additionally, this effect was more pronounced for the modified polymers, particularly 10KS and 10KN, than for PNIPAM-MAA. Further, mixing with the dye reduced negative charges for 10KS and 10KN, but increased negative charges for 10K, 60K, 60KS, and 60KN. When using the dye of different MW, namely 10k dye and 70k dye, we observed similar effects on the size, PDI, and ζ-potential for the 60 kDa polymers.

To ensure DSS-treated mice with more than 15% weight loss, the adhesion experiments were conducted between Day 7 and Day 11 after DSS treatment when the mice with colitis reached the body weight loss% of −15% (Fig. 4A). The body weight loss% of individual mice was monitored, and the colon length on the sacrifice day was measured (fig. S16). The polymers were administered rectally, mice were sacrificed at 24h post-enema, and the dissected colon tissue was for IVIS imaging (fig. S17) and further processing. For each colon tissue, the fluorescence of homogenized colon sections was compared, showing that the middle and distal colon sections possessed higher fluorescence intensity than the proximal colon (fig. S18A). Both the distal and the middle colon also demonstrated a significantly higher MPO activity than the proximal colon (fig. S18B, P < 0.0001), confirming the heterogeneity of the MPO activity in different colon sections and highlighting this biological variation in DSS-induced colitis in mice.

**Fig. 4.**
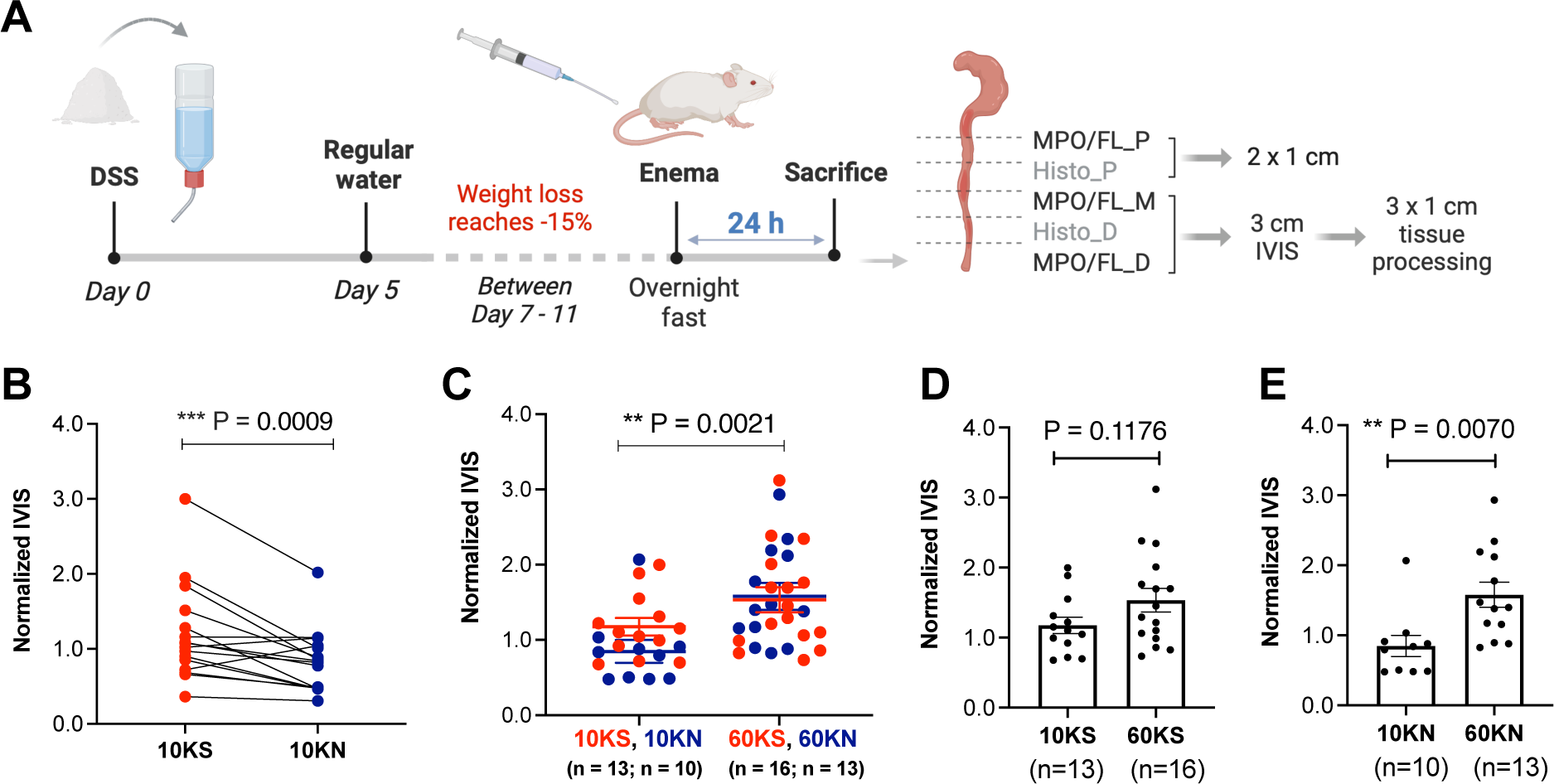
Comparison of polymers’ adhesion to the inflamed colon in DSS-treated mice reaching 15% weight loss. **(A)** Schematic outlining the *in vivo* experimental procedure for mice reaching weight loss of 15%. P: proximal, M: middle, D: distal. **(B)** Paired comparison of similar weight loss% of 10KS and 10KN (n = 16, *P* = 0.0009 was determined by Ratio paired *t* test). **(C)** Comparison of IVIS fluorescence for 10KS, 10KN, and 60KS, 60KN. *P* = 0.0021 was determined by Student’s *t* test. **(D)** There was no significant difference in the fluorescence retention for 10KS and 60KS (*P* = 0.1176 determined by Student’s *t* test). **(E)** There was a significant difference in fluorescence retention between 10KN and 60KN (*P* = 0.0070 determined by Student’s *t* test). Normalized IVIS was calculated by the IVIS fluorescence in the polymer groups divided by the IVIS fluorescence in the dye alone group.

We compared the adhesion between 10KS and 10KN using the 10k dye alone as a control, and the adhesion between 60KS and 60KN using the 70k dye as a control. Here we used normalized fluorescence for comparison in which the fluorescence from the polymer was divided by the fluorescence from the dye alone. We pooled the mice by pairing their weight loss%, using around 15% in this evaluation and our first evaluation from 5% to 20% (Fig. 4B). We found significantly higher fluorescence retention in the inflamed colon for 10KS than 10KN (*P* = 0.0009, ratio paired *t* test for comparable weight loss%). However, we did not observe a significant difference between 60KS and 60KN. The MW of the polymers may play a dominant role in this process than the charges on polymers. With the weight loss% at around 15%, 60KS and 60KN showed significantly higher fluorescence retention than the 10KS and 10KN groups (Fig. 4C, *P* = 0.0021, Student’s *t* test). When we compared the 10KS vs. 60KS, there was no significant difference in the fluorescence retention (*P* = 0.1176) (Fig. 4D); however, 60KN showed significantly higher fluorescence retention than 10KN (*P* = 0.0070) (Fig. 4E). Further, there was no significant difference between the 10k dye and the 70k dye for the adhesion of 60KS (*P* = 0.9177) or 60KN (*P* = 0.8834) to the inflamed colon (fig. S18C, D). Detailed data are tabulated in Tables S1 and S2. The correlations between fluorescence intensity and colitis parameters underscored the heterogeneity in DSS-induced colitis and suggested using comparable colitis parameters to evaluate materials capacity in targeting the inflamed colon.

## Discussion

There is a pressing need to improve current therapies for IBD treatment. Therapeutics that can target the site of inflammation stand to maximize drug efficacy and minimize side effects. Drug delivery systems have employed various mechanisms to target the inflamed mucosa in IBD (*36*). Incorporating more than one targeting strategy in a delivery system has the potential to combine interactions for enhanced localization to the inflamed mucosa. Previous studies, including our work, showed that negatively charged drug delivery systems preferentially targeted the inflamed colon in IBD, owing to the accumulation of positively charged proteins at the site of inflammation (*8, 37*). To increase negative charges on the delivery system and integrate responsiveness to clinically relevant stimuli, we focus on thermo-responsive polymers that provide repetitive units for chemical modification and gel-forming capacity in response to temperature change. Here, we hypothesize that the functionalization of PNIPAM-based polymers can harness the charge-mediated interaction and sol-to-gel transition for enhanced targeting to the inflamed colon in experimental models of colitis. We characterized these microgel-based soft colloidal systems and investigated these polymers interacting with the inflamed colon, which could bring new approaches to target the site of inflammation for IBD treatment.

In this work, PNIPAM-MAA was chemically modified with taurine (S modification) to increase negative charges and with *N,N*-dimethyl-ethylenediamine (N modification) to decrease negative charges. The reactions were conducted for PNIPAM-MAA of two molecular weights, 10 and 60 kDa, respectively. The resultant modified polymers were 10KN, 10KS, 60KN, and 60KS. The unmodified polymers, 10K and 60K, were as a control. The ζ-potential of these thermo-responsive polymer-based microgels confirmed the structure modification: 60KS and 10KS with the most negative charges, 60K and 10K being less negative, and 60KN and 10KN being the least negative. For these polymers alone, we noticed that the ζ-potential increased as the temperature increased from 25°C to 37°C. From a colloid chemical point of view, this increase in the ζ-potential stabilized the microgels by counteracting the instability usually enhanced at higher temperatures (*38*).

The chemical modification not only changed the surface charge but also altered the hydrophobicity/hydrophilicity of polymers, thereby affecting the LCST. The sulfonates in the S modification are strongly hydrated anions, while the tertiary amines in the N modification are weakly hydrated cations. Both ions are important species in the Hofmeister ion series that exert specific ion effects on the water solubility of macromolecules and the behavior of colloidal systems in aqueous solutions (*39, 40*). When present as ions in solutions, the sulfonates and tertiary amines contribute strongly to the “salt-out” process that tends to stabilize macromolecules and reduce the LCST. However, such an effect on the LCST would be different when the ligand was conjugated to macromolecules, instead of being present as salts in the solution. It has been reported that hydrophilic dextran increased the LCST of PNIPAM-based polymers when grafted to the polymer and reduced the polymer’s LCST when present in the solution, exerting opposite effects on the polymer’s thermo-responsiveness (*41*). In our work, the primary effect of the grafted sulfonates and tertiary amines on PNIPAM-MAA was to increase the hydrophilicity of the polymer, thereby increasing the LCST. This was consistent with the fact that the modified polymers required a higher concentration than PNIPAM-MAA to achieve *T*_cp_ = 37°C. Furthermore, due to the presence of sulfonates or tertiary amines in the polymers, they likely contributed to stabilizing the polymeric networks and forming microgels.

Because the LCST of PNIPAM-based polymers is related to their concentrations and molecular weights (*42*) that directly affect the gelation process, one unique challenge in our work was how to compare these polymers for *in vivo* studies to determine the charge effect on the polymers’ targeting to the inflamed colon. We chose to compare them at individual polymers’ CGC of *T*_cp_ = 37°C to achieve the *in-situ* gelation after rectal administration. Owing to their different chemical compositions and molecular weights, 10K, 10KN, 10KS, 60K, 60KN, and 60KS were used at 5.6, 32.4, 38.7, 4.9, 10.1, and 16.5 mg/ml for *in vivo* experiments. The corresponding concentration of the 60K-based polymers was lower than the 10K-based polymers, likely due to the increased polymer chain length, thereby increasing the opportunities to undergo hydrogen bonding to form microgels.

DSS-induced acute colitis was chosen due to the rapid onset of inflammation and the well-established procedure for generating this model. To minimize the heterogeneity in experiments, we removed mice that did not lose weight after the 5-day DSS treatment on day 7. Further, weight loss% was stratified to ensure appropriate within-group comparisons among animals and comparable average weight loss% for each group. Owing to the self-recovery nature and limited disease duration of this acute model (*28*), we determined the polymers’ adhesion before the full recovery of colitis, either on Day 7 for 6 h post-enema evaluation or by Day 11 for 24 h post-enema evaluation when using the weight loss of around 15%. We found that for 10K-based polymers, the charge interactions dominated the binding and led to higher fluorescence retention for 10KS than 10KN, as shown by the ratio of Inflamed/Healthy and the ratio paired *t*-test using similar weight loss%. In contrast, for 60K-based polymers, the charge-mediated interaction was not dominant. Although 60KN showed the least negative charge, it exhibited comparable fluorescence retention in the inflamed colon to 60KS and 60K. All 60K-based polymers retained higher fluorescence in the inflamed colon than the 10K-based polymers, indicating the importance of molecular weight in the adhesion process.

An important feature of this study is that we examined the correlation between polymer targeting and colitis parameters in individual mice from the DSS-induced acute colitis and healthy controls. The heterogeneous nature of chemically induced colitis is well-known, but how the non-uniformity of the chemically induced colitis may interfere with the comparison of polymers targeting the inflamed colon has been underappreciated. We analyzed the colitis parameters, including colon length, weight loss%, MPO activity, and histology score, and found that, in general, the polymer targeting was aligned with colitis severity. The higher the histology scores and the MPO activity and the shorter the colon length, the higher the fluorescence retention in the inflamed colon; although we did not observe a statistically significant correlation in our analysis for these three parameters. On the other hand, the weight loss% correlated significantly with the polymers’ targeting. Within each polymer group, we observed that the higher the weight loss%, the higher the polymer binding to the inflamed colon. The PCA analysis also confirmed the significant contribution of weight loss to fluorescence, indicated by a significantly nonzero contribution using the least squares regression (p < 0.0001). This led to our study using colitic mice with similar weight loss% to compare the charge effect on polymers’ targeting for 10KS and 10KN. As a result, we found significantly higher fluorescence retention by 10KS than by 10KN.

The interaction of polymers with the inflamed mucosa *in vivo* is complex and could be a combination of several factors, including binding to the accumulation of the positively charged proteins locally, the increased intestinal tissue permeability, the sol-to-gel phase transition, and the microgel particles internalized by the infiltrated immune cells. Given this complex context, our study supports using mice of similar weight loss% in chemically induced colitis models when comparing materials’ binding capacity to the inflamed colon. Further, the weight loss% can be monitored daily and measured routinely for assessment, compared to other colitis parameters that can only be analyzed after euthanization.

With regards to *in vivo* tracking, we chose to mix a fluorescent dye, Texas Red^®^ dextran, with the polymer to form microgels. Although chemically conjugating a dye to the polymer has the advantage of monitoring the microgel long-term, it bears the risk of altering the polymer structure and the LCST, thereby interfering with the polymer behavior *in vivo*. As a hydrophilic polysaccharide, dextran was known to improve water retention and drug release profile for PNIPAM-based microgels by reducing the demixing/syneresis phenomenon (*41*). Therefore, in addition to tracking the polymer *in vivo*, using a Texas Red^®^ dextran dye could also provide information on the effect of additives for PNIPAM-based materials for future experiments. We also examined the effect of the concentration and molecular weight of the Texas Red^®^ dextran on the size and ζ-potential of the microgels. We found that mixing with the dye reduced the particle size and PDI of polymer microgels as the dye concentration increased for all polymers. The effect on the ζ-potential varied, depending on the dye’s molecular weight and concentration. However, adding the dye did not change the pattern of the size and ζ-potential for all polymers, compared to individual polymers without mixing with the dye. At 37°C, the dye alone formed nanocomplexes of about 200 nm with negative charges, consistent with previous reports on dextran nanoparticles (*43*). These negatively charged dextran nanocomplexes could explain our observation that the dye alone showed higher fluorescence retention in the inflamed colon than the healthy colon, a combination of charge-mediated interaction and passive deposition due to the increased intestinal permeability in colitic mice. In our experiments, the dye-alone group, including 10k dye and 70k dye, was studied in parallel with all polymers. When comparing polymers, the fluorescence from each polymer group was normalized against the corresponding dye-alone group.

Since the chemical modification not only changed the surface charge but also affected the interactions between the chemical groups in the polymer, limitations on characterization remain for the TRIT delivery systems based on the PNIPAM polymers. Both 10KS and 60KS are optimal candidates for the TRIT system, given their high fluorescence retention to the inflamed colon than the healthy colon; further, the grafted taurine will provide the well-established beneficial biological effects (*23–27*). Due to the difference in polymer molecular weights, suitable applications for each polymer need to be determined individually. Unlike well-established colloidal “hard sphere” models, thermo-responsive microgels are soft colloidal systems and far more versatile with complex behaviors (*44*). Multiple factors that affect their thermo-responsiveness, including the polymer structure, density, ionic strength, and the microgel’s impact on the local dynamics of the dispersion, have still to be explored. Additionally, PNIPAM-MAA is not only thermo-responsive but also pH-responsive. The measurement of the colon content in normal mice showed a pH of 7.8 and in mice with colitis a pH of 6.7 (*45*). The lower value in colitis than the normal mice was consistent with reports in the human IBD (*46*), although the degree of acidity may vary depending on disease severity. The more acidic, the less negatively charged the carboxylic acids are since they are weak acids. However, it will unlikely affect the strong sulfonic acids from taurine. In this work, we did not evaluate the pH effect on the gelation, because we considered that the negative charge from the sulfonates will unlikely be affected by the physiological pH, even under inflammation. Another factor that may affect the extrapolation and translation of these results to human studies is the thermoregulatory difference in the physiological temperature of mice and humans (*47*). Mice tend to have a widespread and on average slightly lower core temperature than humans. Therefore, using the polymer concentration of *T*_cp_ = 37°C that we reported here may achieve a more complete gelation in humans than in mice. Further studies on the physicochemical properties of PNIPAM-MAA and the modified polymers may inform additional implications when using thermo-responsive microgels in a broad area of biological and biomedical applications, including designing polymer-based drug delivery systems.

In sum, we report an approach to chemically modify thermo-responsive PNIPAM-based materials for targeted drug delivery to the inflamed colon in experimental colitis using charge-mediated interaction combined with the sol-to-gel transition. The charge-mediated interaction was dominant at a low molecular weight of 10 kDa but not for polymers at a high molecular weight of 60 kDa, despite the chemical modifications. The correlation between polymer targeting and colitis parameters suggested using similar weight loss% when comparing materials interaction in DSS-induced acute colitis. Further understanding of polymeric hydrogels interacting with the inflamed intestinal mucosa could refine the future design of drug delivery systems in IBD. Here, our study to characterize and analyze PNIPAM-based microgels provides a delivery system that can adhere specifically to the inflamed colon, a potential platform for maximal efficacy and minimal side effects in colonic IBD treatment.

## Materials and Methods

### Materials

Taurine, N,N-dimethyl-ethylenediamine, N-hydroxy-sulfosuccinimide (Sulfo-NHS), 1-ethyl-3-(3-dimethyl-aminopropyl)carbodiimide hydrochloride (EDC), 2-(N-morpholino) ethane-sulfonic acid) (MES), Hexadecyltrimethyl ammonium bromide, O-dianisidine dihydrochloride, hydrogen peroxide (stabilized, 30 wt.% in H_2_O), and Poly(N-isopropylacrylamide-co-methacrylic acid) (10 mol% in methacrylic acid, PNIPAM-MAA) with a molecular weight of 10k and 60k, respectively, were purchased from Sigma Aldrich, Inc. Dextran Texas Red™ (Neutral, MW = 10K and 70K) was purchased from ThermoFisher Scientific, Inc. Dextran sulfate sodium (DSS) was purchased from Affymetrix, Inc. Due to a supply shortage from Sigma Inc., part of the 10k PNIPAM-MAA was synthesized in-house (*21*). For polymer synthesis, N-isopropylacrylamide (NIPAM), methacrylic acid (MAA), and azobisisobutyronitrile (AIBN) were purchased from Sigma Aldrich. The in-house synthesized PNIPAM-MAA was confirmed by ^1^H-NMR. The MW was determined by Gel Permeation Chromatography (GPC, water phase) and compared with the purchased one, indicating comparable *M*_n_ and polydispersity (*M*_w_/*M*_n_).

### Synthesis and characterization of modified PNIPAM-MAA

#### Modification of PNIPAM-MAA with S and N ligands

PNIPAM-MAA was modified with S or N ligands via the EDC/NHS coupling reaction. The molar ratio of 1:3:3 between reactants (methacrylic acid: EDC: sulfo-NHS) was used. First, 100 mg of 10k or 60k PNIPAM-MAA (0.0905 mmol of methacrylic acid) was dissolved in 6mL of MES buffer (0.1 M, pH = 5.2) until completely dissolved. After six hours, sulfo-NHS (58.98 mg, 0.272 mmol) was dissolved in 1 ml of MES buffer. Then 1ml of sulfo-NHS and EDC (59.37 µl, 0.272 mmol) were quickly added to the polymer solution, and the mixture was kept on a shaker for 45 mins. For S modification, 113.3 mg of taurine (0.905 mmol) dissolved in 7ml of PBS (10x) was added and kept on a shaker overnight. For N modification, the entire reaction was repeated using 98.9 µl of N, N-dimethyl-ethylenediamine (0.905 mmol) instead of taurine. Taurine (S modification) and N,N-dimethyl-ethylenediamine (N modification) were added at a 1:10 molar ratio of the MAA in the polymer. After overnight shaking, polymer solutions were dialyzed for 4 days with dialyzed against carbonate buffer (0.2 M, pH = 10) (x4) first and then dialyzed against water (x3). The dialysis tubing’s molecular weight cutoff (MWCO) was 1.0 kDa for the 10K polymers and 3.5 kDa for the 60K polymers. The polymers were then lyophilized and stored for future testing. The chemical structure of modified polymers was confirmed by ^1^H-NMR. In a typical synthesis procedure for the 10k PNIPAM-MAA, NIPAM (916.6 mg, 8.1 mmol), AIBN (44.3 mg, 0.27 mmol), methacrylic acid (MAA) (77.8 mg, 0.9 mmol), and anhydrous dimethylformamide (DMF, 6mL in total) were added to a 25 ml Schlenk round bottom flask. The mixture was stirred for 15 min to dissolve fully. Then the solution was degassed by purging with nitrogen for another 15 min. Polymerization was carried out in an oil bath at 85°C and stirred for 16 hours. After polymerization, the mixture was precipitated by adding dropwise into pre-chilled diethyl ether under vigorous stirring (200 ml, 3x). The resultant white solid was dried under vacuum overnight, dialyzed (MWCO 3.5 KDa) against water for seven days to remove residual DMF, and then lyophilized to provide 664.5 mg of polymer (yield, 66.8%). The synthesized polymers were analyzed by ^1^H-NMR and water-phase gel permeation chromatography (GPC). The GPC analysis of the 10k PNIPAM-MAA from Sigma showed *M*_n_ of 9,115 Daltons and *M*_w_/*M*_n_ = 2.876, and the three batches of in-house synthesized 10k PNIPAM-MAA showed *M*_n_ of 10,313 Daltons with *M*_w_/*M*_n_ = 2.819, *M*_n_ of 9,726 Daltons with *M*_w_/*M*_n_ = 2.022, and *M*_n_ of 11,066 Daltons with *M*_w_/*M*_n_ = 2.349.

#### LCST determination using the Cloud Point method

To determine the lower critical solution temperature (LCST), PNIPAM-MAA or modified polymers were dissolved in 1× PBS for a range of concentrations (fig. S2). The change in the absorbance as a function of temperature was recorded by a Cary 100 Bio UV-vis spectrophotometer (Agilent Inc.). The temperature ranged from 25°C to 50°C, with a heating rate of 0.50°C/min and a data collection interval of 0.5°C. The absorbance was measured at a wavelength of 420 nm and then converted to transmission% (T%) (*48*). The LCST was determined to be the inflection point in the transmittance curve. It was calculated as the temperature at which the magnitude of change in transmittance over the temperature change was maximal by determining the maximal value of the first derivative of the transmittance curve (*21*). The four-parameter logistic regression was performed on the polymer concentration vs. LCST data to interpolate the concentration value that would yield the *T*_cp_ = 37°C. Then the absorbance of these polymer concentrations interpolated from the fitted curve was measured using the Could Point method across the temperature range 25°C to 50°C. These concentration values were confirmed experimentally to show that the LCST of individual polymers were between 36.5 and 37.5 °C.

#### Size and zeta potential (ζ-potential) measurement

The hydrodynamic size *D*_h_ (Z-average intensity mean) and ζ-potential of the polymers were measured by a NanoSeries Malvern. For measurements across a temperature range, the initial temperature was 25°C and was then increased at a 2°C interval to 50°C. Each measurement had a 2-minute temperature equilibration time. The concentration of the individual polymers for the temperature ramp measurements was 0.5 mg/mL, and the dispersant was 1mM NaCl. Each sample was measured at each temperature point three times, and this entire cycle from 25°C to 50°C was repeated three times for each polymer. Size and ζ-potential were also measured at 25°C and 37°C, respectively, with a 5-minute temperature equilibration time. The polymers were measured at individual concentrations of *T*_cp_ = 37°C. The dispersant was PBS (1×, pH = 7.4). Each polymer was measured three times at each temperature.

#### Cryo-scanning electron microscopy (SEM)

To ensure complete gelation at 37°C for imaging, we prepared each modified polymer 60KS and 60KN in water at 2-fold of their CGC at *T*_cp_ of 37°C. The concentrations for individual polymers were 33 mg/mL of 60KS and 20.2 mg/mL of 60KN. The polymer solutions were heated to gelation, snap frozen, coated with 80:20 Pt/Pd, and imaged. Briefly, one drop of the sample was preheated to 50°C, then immediately frozen in nitrogen slush at −220°C. Then the sample was cryo-transferred under a vacuum in the cryo-fracture apparatus chamber (Baltec MED-020), where it was fractured at −145°C. The temperature was then decreased to −100 °C and maintained at this temperature for 10 min for sublimation. The sample was then coated with 80:20 Pt/Pd for 100 s (∼10nm thickness) and introduced into the microscope chamber for imaging. Cryo-SEM images were obtained using the Zeiss NVision 40 microscope, operating at a 2-3kV acceleration voltage.

### Colitic mice model

Adult BALB/c wild-type (WT) mice were purchased from the Jackson Laboratory. Experiments involving WT mice were performed at the Massachusetts Institute of Technology’s Koch Institute. All mouse studies were performed according to institutional and NIH guidelines for humane animal use. DSS colitis was induced in 7-8-week-old BALB/c mice by feeding them 3.0% DSS in drinking water for 5 consecutive days. Mice were then switched to regular water. The body weight of mice was measured daily. To minimize the heterogeneity in experiments, mice that did not lose weight after the 5-day DSS treatment were not used for experiments.

### *In vivo* adhesion evaluation of polymers

For *in vivo* adhesion testing, mice with colitis and their disease-free controls (untreated WT) were fed the Alfalfa-free diet (Envigo Inc.) to reduce the autofluorescence background. The body weight of mice was measured daily to monitor the disease development. On Day 6, mice were weighed and then fasted overnight to prepare for enemas the following morning. On Day 7, mice with different weight loss% were regrouped based on Day 6 weights to ensure the average weight loss% was roughly equal in each colitis group for polymer comparison. Then each mouse received an enema of 100 μl of the polymer solution containing Texas Red^®^ Dextran in phosphate-buffered saline (PBSx1, pH = 7.4). Briefly, individual mice were anesthetized with 2.5% isoflurane, a 20G flexible disposable feeding needle (Cadence Science, Inc.) was advanced into the rectum 3 cm past the anus, the polymer solution with dye was administered, the catheter was removed, and the anus was kept closed manually for 1 minute before releasing the mouse into the cage. Animals were sacrificed after 6 hours. The distal 3 cm of the colon was removed and imaged using an IVIS fluorescence imager (IVIS 200, Perkin Elmer). Fluorescent signal intensity was quantified using Living Image 4.3.2 software (IVIS 200, Perkin Elmer). In another set of experiments, the adhesion between polymers was compared using only colitic mice with weight loss% of 15 - 20%. Mice were fed drinking water containing 3% DSS for 5 days and then switched to regular drinking water. Starting from Day 7, only mice with a weight loss percentage between 15 - 20% were used for the adhesion experiment. Then the mice were sacrificed at 24 h after administering the polymer and dye mixture. Owing to this acute model’s self-recovery nature and limited disease duration, we evaluated the polymers’ targeting by Day 11 before the full recovery of colitis.

### Myeloperoxidase (MPO) activity

The distal colon was homogenized in 1:20 (w/v) of 50 mM phosphate buffer (pH = 6) containing 0.5% hexadecyltrimethyl ammonium bromide on ice using a homogenizer (Bertin Corp., Precellys).

The homogenate was centrifuged at 14,000 rpm for 15 minutes. The supernatant (10 μl) was added to 190 μl of 50 mM phosphate buffer (pH 6) containing 0.167 mg/ml O-dianisidine dihydrochloride and 0.0005% hydrogen peroxide. The change in absorbance at 460 nm was measured. One unit of MPO activity is defined as degrading 1 μmol of hydrogen peroxide per minute at 25°C (*8*). Total protein content was determined by BCA protein assay (ThermoFisher Scientific Inc.).

### Histology analysis

All mice were sacrificed for histopathological analysis on day 7 after IVIS imaging. Colons were isolated, fixed in 4% Paraformaldehyde, and embedded in paraffin. Standard Hematoxylin and Eosin (H&E) - stained sections were examined and scored by an experienced pathologist (J.K.L.) in a blinded fashion. The parameters mononuclear cell infiltration, polymorphonuclear cell infiltration, epithelial hyperplasia, and epithelial injury were scored as absent (0), mild (1), moderate (2), or severe (3), giving a total score of 0-12.

### Statistical analysis

Statistical analysis and graphing were performed with Prism version 10.0.2 (GraphPad Software). The two-tailed Student’s *t* test was used to compare differences between two experimental groups, except for the study comparing polymers for targeting inflamed colon using similar weight loss%, where the Ratio paired *t* test was used. A value of *p* < 0.05 was considered statistically significant.

## Acknowledgments

We thank the Koch Institute Swanson Biotechnology Center at the Massachusetts Institute of Technology for their technical support, specifically the Preclinical Imaging & Testing Core and the Histology Core. We also thank S. Afewerki for assistance with the in-house synthesis of 10K PNIPAM-MAA. The schematics in Figs. 3A and 4A were created with Biorender.

## Funding

S.Z. acknowledges funding from the National Institutes of Health (NIH; R01 DK136941), American Gastroenterological Association (AGA, AGA-Pfizer Pilot Research Award in Inflammatory Bowel Disease AGA202021-21-06), and the Crohn’s and Colitis Foundation (Litwin IBD Pioneers Award # 993684). In addition, this work was supported by a KL2/Catalyst Medical Research Investigator Training Award (CMeRIT Award to S.Z.) from Harvard Catalyst | The Harvard Clinical and Translational Science Center and financial contributions from Harvard University and its affiliated academic healthcare centers. The content is solely the responsibility of the authors and does not necessarily represent the official views of Harvard Catalyst, Harvard University and its affiliated academic healthcare centers.

## Author contributions

S.Z. and A.T.J. designed and performed the experiments, analyzed the data, and wrote the manuscript. W.T., R.Z., L.J., Y.Z., and H.Z. contributed to collecting experimental data, helped analyze the data, and edited the manuscript. J.K.L. analyzed histology samples. J.R.K., R.L., and G.T. reviewed and edited the manuscript.

## Competing interests

S.Z., J.R.K., R.L., and G.T. are co-inventors on a patent (U.S. Patent No. 11,684,583) encompassing the technology described in this paper. All other authors declare they have no competing interests.

## Data and materials availability

All data are available in the main text or the supplementary materials.

## Supplementary Materials

### This PDF file includes

Figs. S1 to S18

Tables S1 to S2

