## Supplementary Materials for "Thermo-Responsive Polymers Targeting Inflammation in Murine Colitis"

Sufeng Zhang *et al.*

**This PDF file includes:**

Figs. S1 to S18
Tables S1 to S2

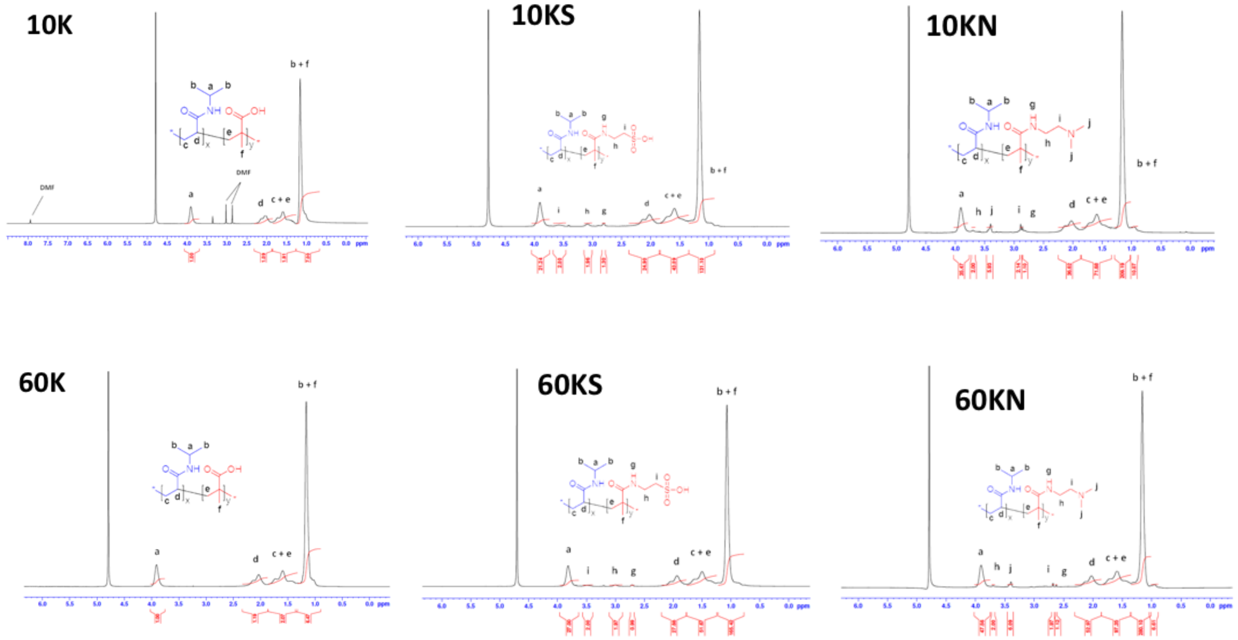


**Fig. S1. NMR spectra of the unmodified and modified PNIPAM-MAA.** The $H^1$-NMR spectrum confirmed the chemical structures of the unmodified and the modified polymers.

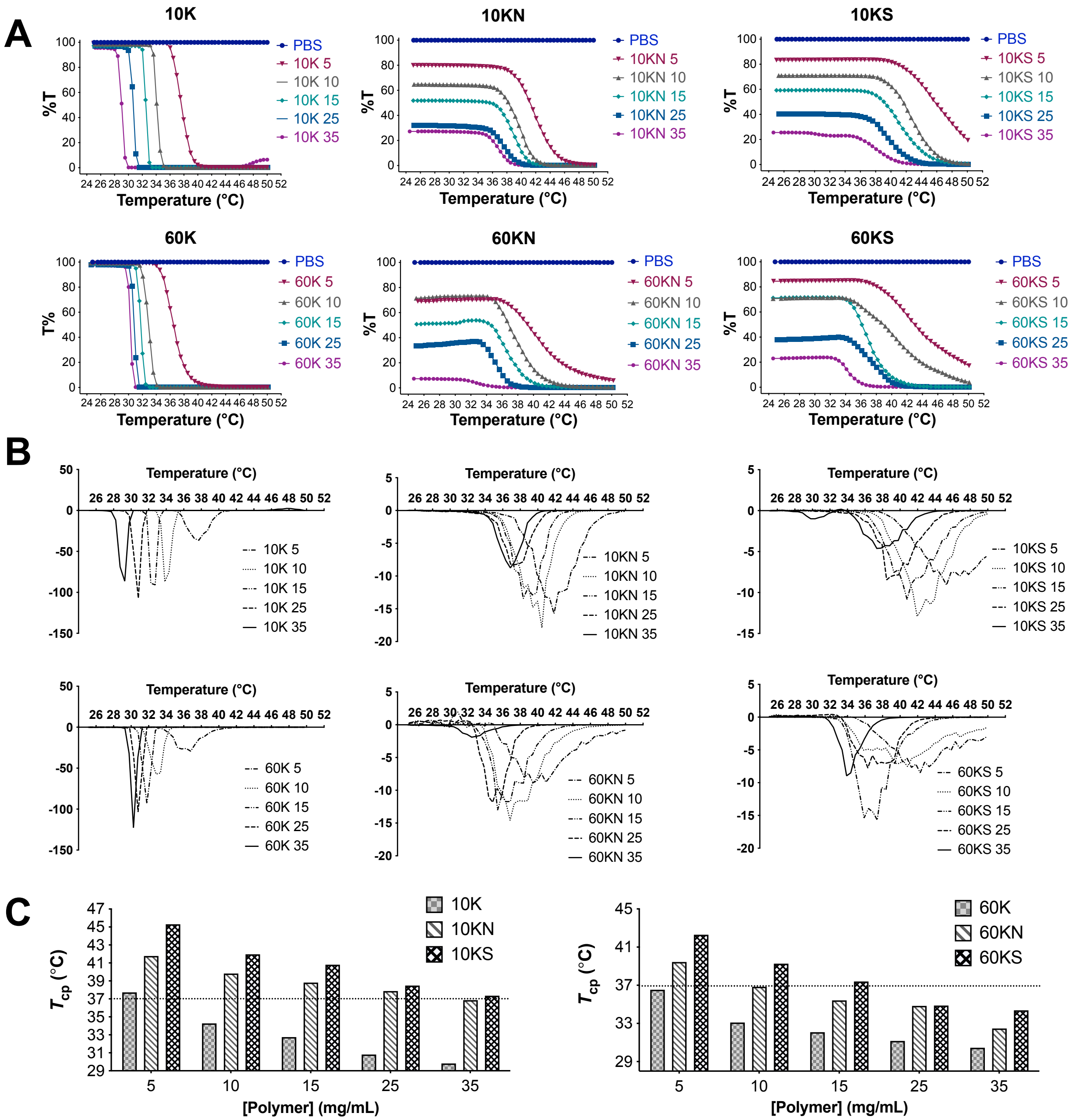


**Fig. S2. Cloud Point ($T_{cp}$) determination for the polymers of different concentrations. (A)** Transmittance% (T%) as a function of temperature for PNIPAM-MAA (10K, 60K) and the modified polymers (10KN, 10KS, 60KN, and 60KS) of different concentrations in PBS. (**B**) First derivatives of the T% curves as a function of temperature for all polymers evaluated at different concentrations for $T_{cp}$ determination. (**C**) $T_{cp}$ values were calculated from the first derivatives of the T% curves vs. temperature for 10K, 10KN, 10KS, 60K, 60KN, and 60KS at different concentrations.

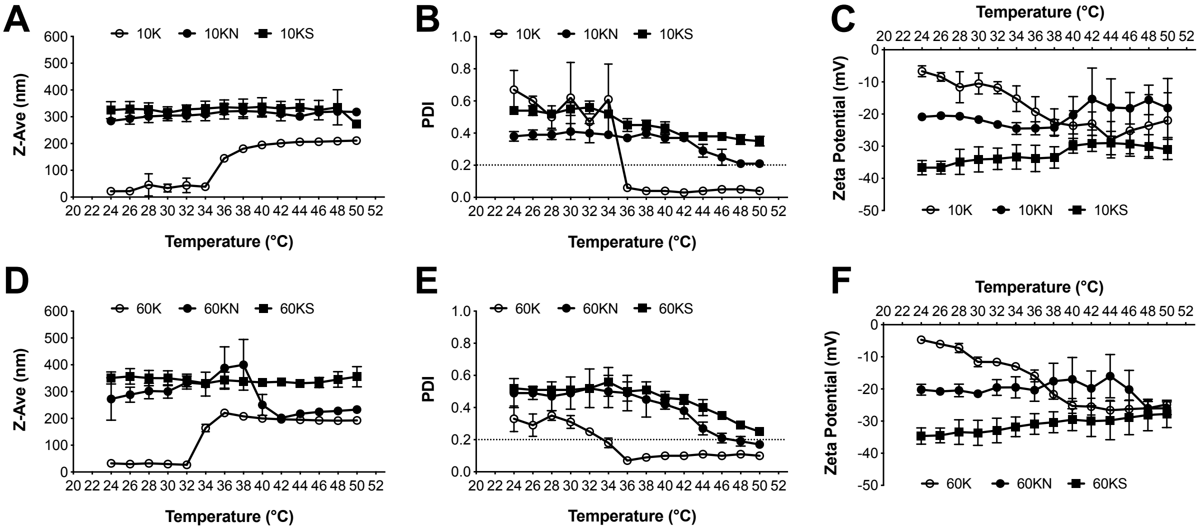


**Fig. S3. Hydrodynamic size and zeta potential of the polymers at 0.5 mg/ml as a function of temperature.** (**A - C**) Size, PDI, and zeta potential measurement for 10K, 10KN, and 10KS at 0.5 mg/ml. (**D - F**) Size, PDI, and zeta potential measurement for 60K, 60KN, and 60KS at 0.5 mg/ml. The temperature ranged from 24°C and was then increased at a 2°C interval to 50°C at a rate of roughly 0.2°C/min.

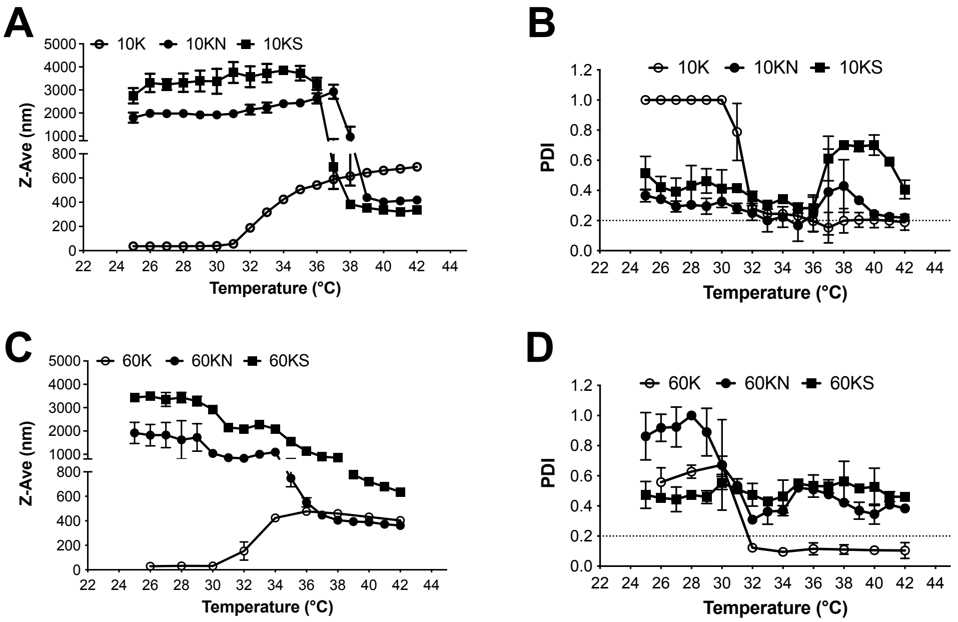


**Fig. S4. Hydrodynamic size of the polymers at the concentrations of $T_{cp}$ = 37°C as a function of temperature. (A)** Size and **(B)** PDI measurement of 10K, 10KN, and 10KS as a function of temperature for the polymers at their concentrations of $T_{cp}$ = 37°C. For 10K, 10KN, and 10KS, the concentration of each polymer at $T_{cp}$ = 37°C is 5.6 mg/ml, 32.4 mg/ml, and 38.7 mg/ml. (**C**) Size and **(D)** PDI measurement of 60K, 60KN, and 60KS as a function of temperature for the polymers at their concentrations of $T_{cp}$ = 37°C. For 60K, 60KN, and 60KS, the concentration of each polymer at $T_{cp}$ = 37°C is 4.9 mg/ml, 10.1 mg/ml, and 16.5 mg/ml. The temperature was evaluated from 25°C and was then increased at a 1°C interval to 42°C for 10K, 10KN, and 10KS; it was measured at a 2°C interval from 26°C to 42°C for 60K, 60KN, and 60KS.

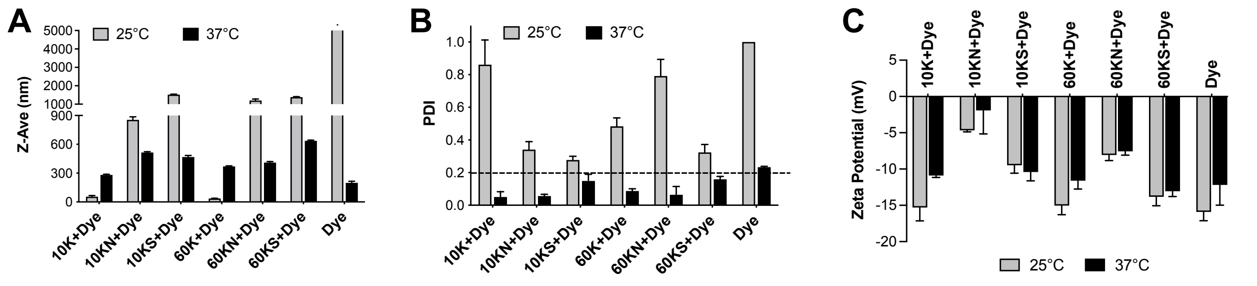


**Fig. S5. Characterization of the polymers at the concentrations of $T_{cp}$ = 37°C with a dye (10k dye, Texas Red® Dextran) for *in vivo* studies.** These polymers were evaluated by (**A**) size, (**B**) PDI, and (**C**) zeta potential measurement. The dye concentration used for the animal experiment was 0.1 mg/ml. The size, PDI, and zeta potential were evaluated at 25°C and 37°C, respectively.

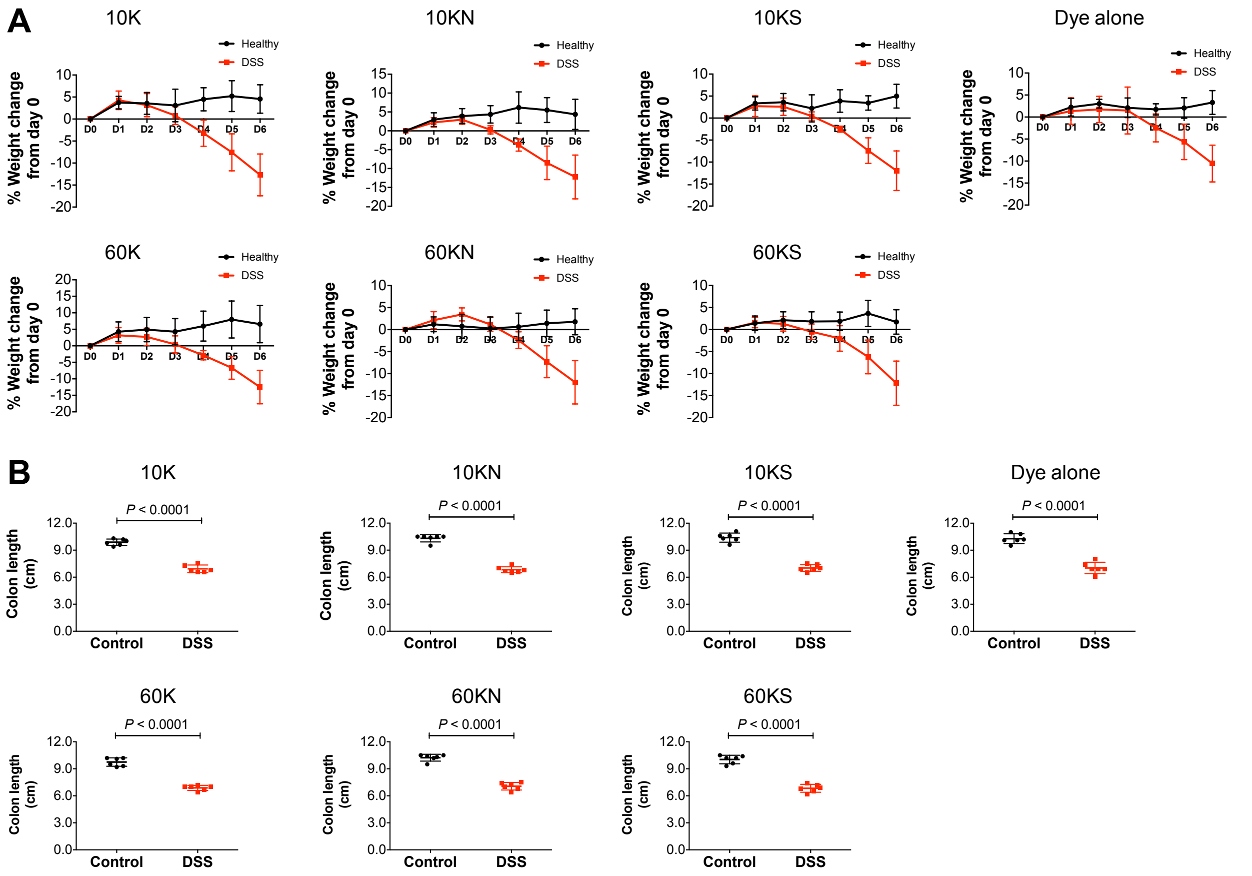


**Fig. S6. Body weight loss% and colon length for the *in vivo* evaluation. (A)** The weight loss% of mice used for polymer evaluation *in vivo*. **(B)** The colon length of mice used for polymer evaluation *in vivo*.

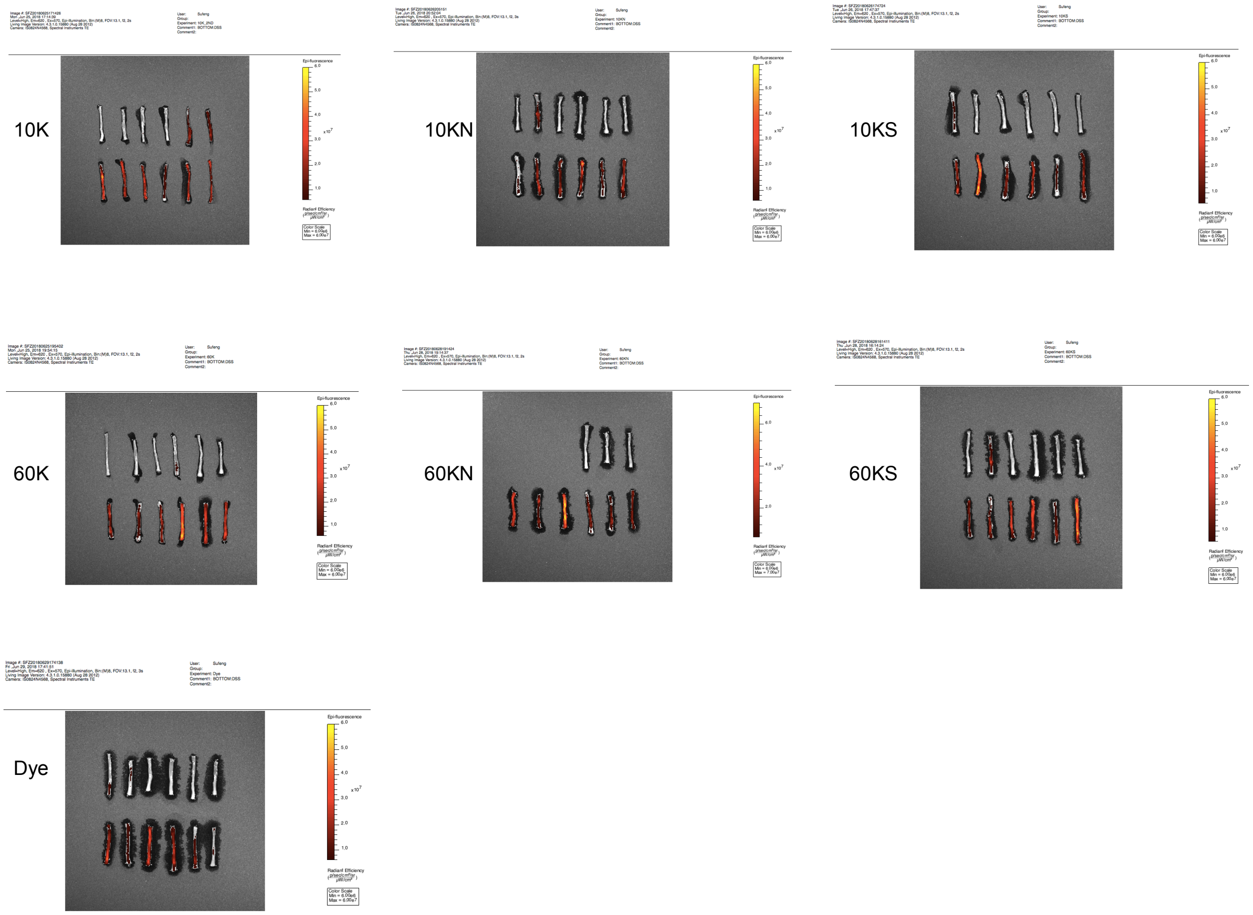


**Fig. S7. IVIS imaging of colon tissue harvested from mice for *in vivo* evaluation.** The polymers used were 10K, 10KN, 10KS, 60K, 60KN, 60KS, and the dye alone.

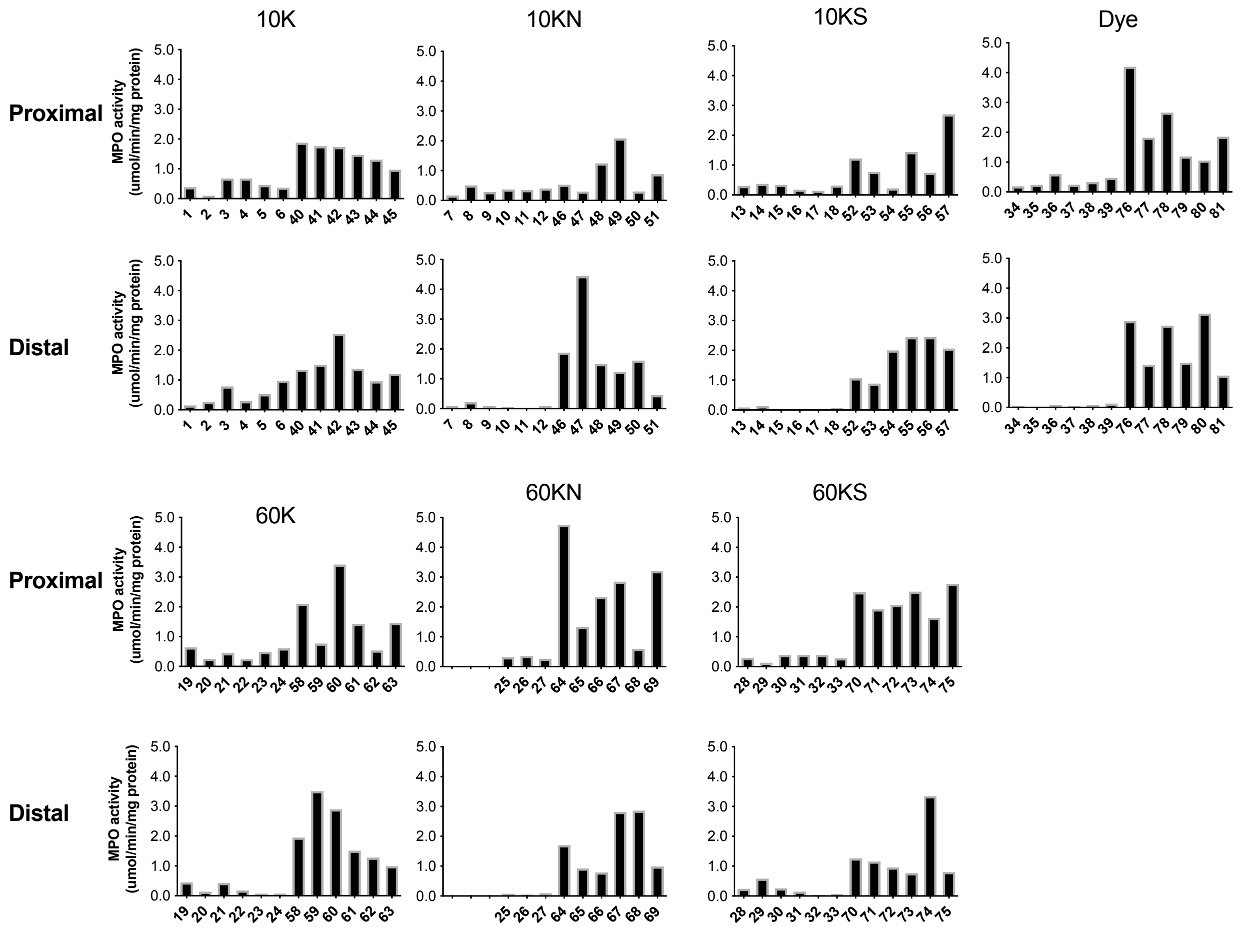


**Fig. S8. MPO activity of the proximal and distal colon for individual mice.** Two pieces of colon tissues collected from individual mice after the IVIS imaging were used for MPO analysis. The x-axis indicates individual mice from #1 - #81. The MPO results of the dissected tissue showed the heterogeneity of MPO activity across the colon tissue in the same mouse and across individual mice in the group.

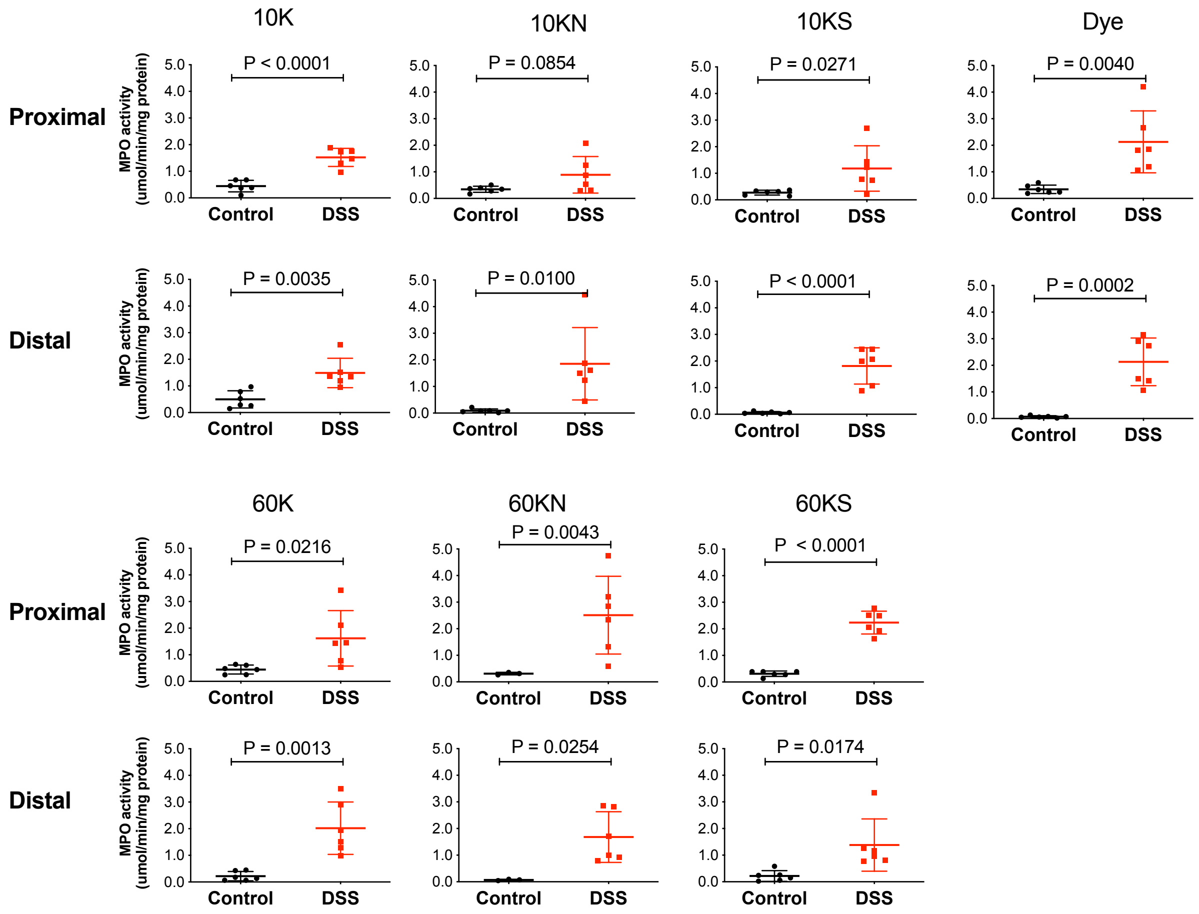


**Fig. S9. Averaged MPO activity of mice in the Control and DSS-treated groups.** The analysis confirmed that mice treated with DSS showed significantly higher MPO activity than the healthy controls. Both the proximal and the distal colon pieces were evaluated.

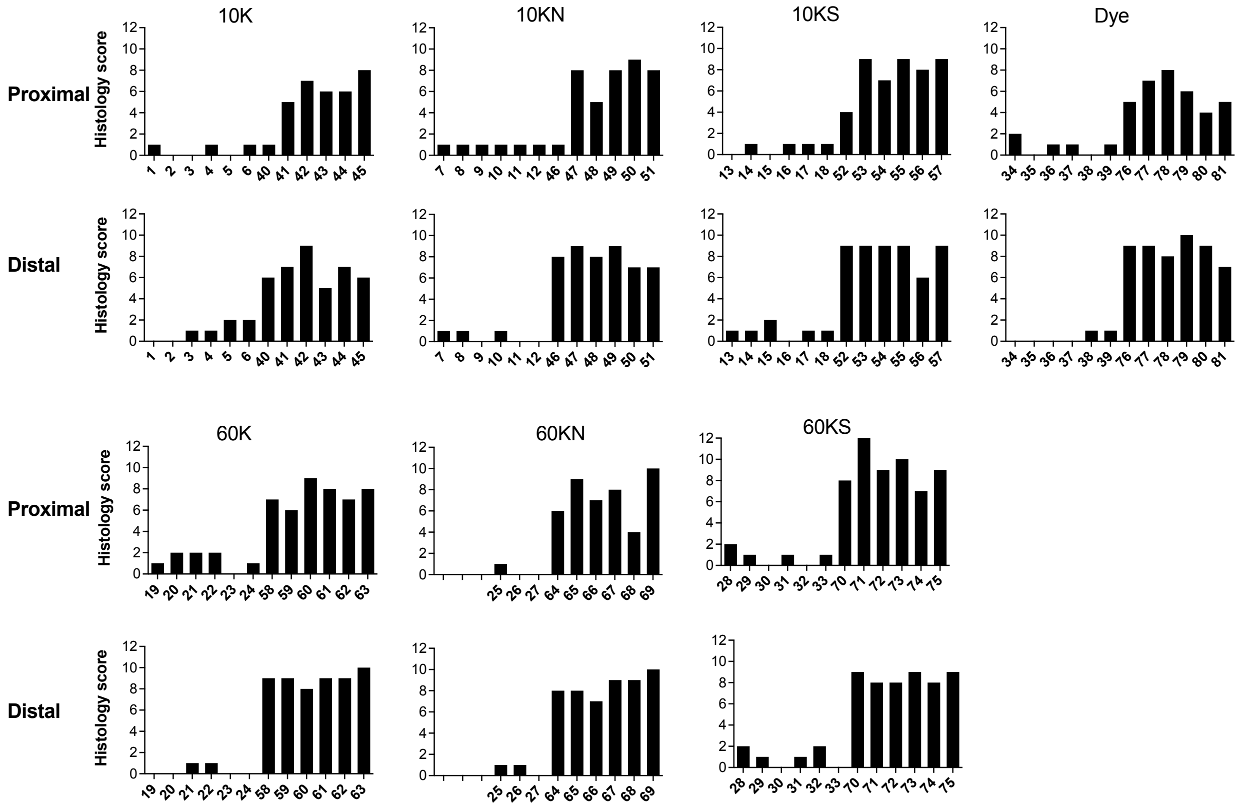


**Fig. S10. Histology analysis of the proximal and distal colon tissues for individual mice.** After the IVIS imaging, two pieces of tissue were collected from each mouse. The x-axis indicates individual mice from #1 - #81. Analysis of the dissected tissue showed the heterogeneity of histology across the colon tissue in the same mouse and across individual mice in the groups.

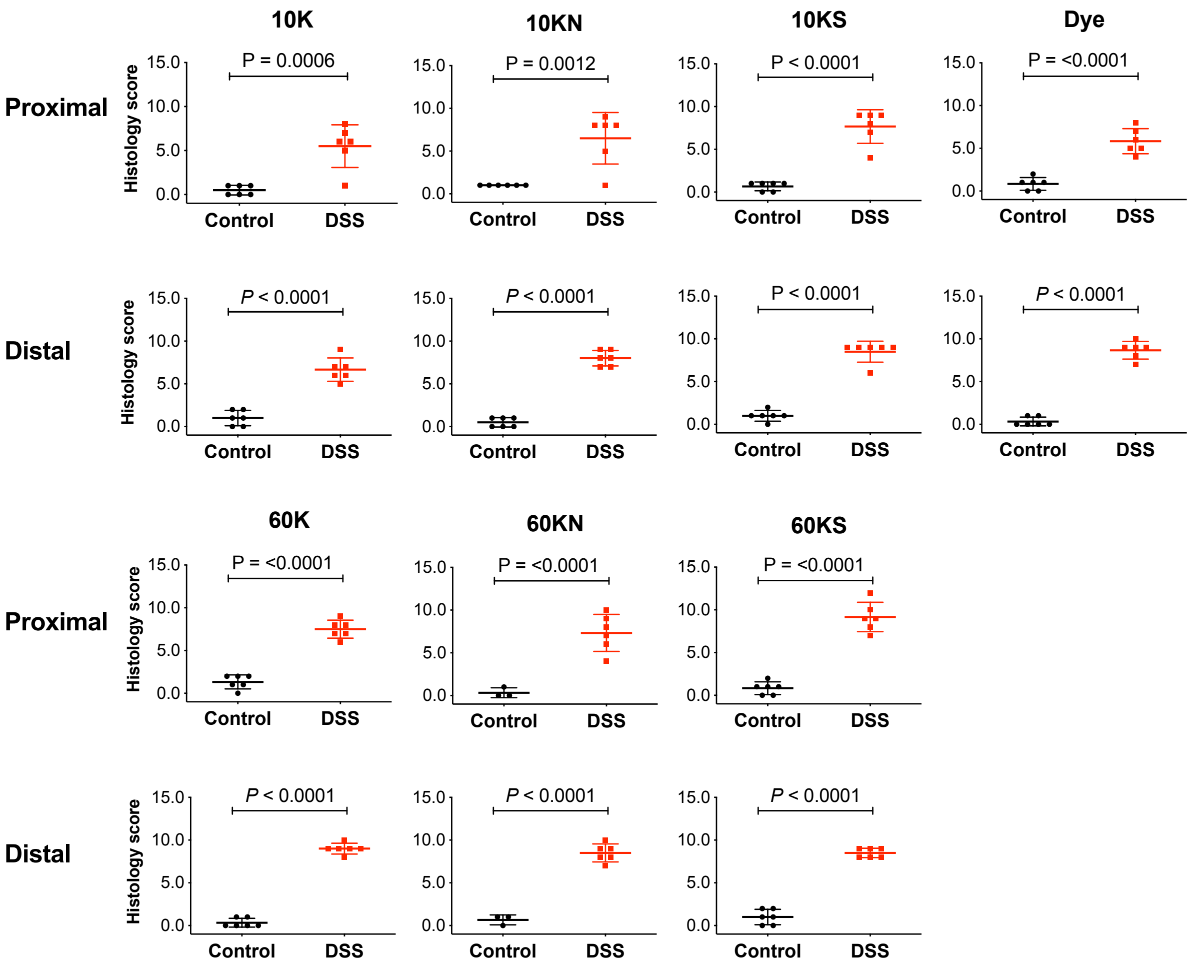


**Fig. S11. Averaged histology analysis of mice in the Control and DSS-treated groups.** The results confirmed that the DSS-treated mice showed significantly higher histology scores than the healthy controls. Both the proximal and the distal colon pieces were evaluated.

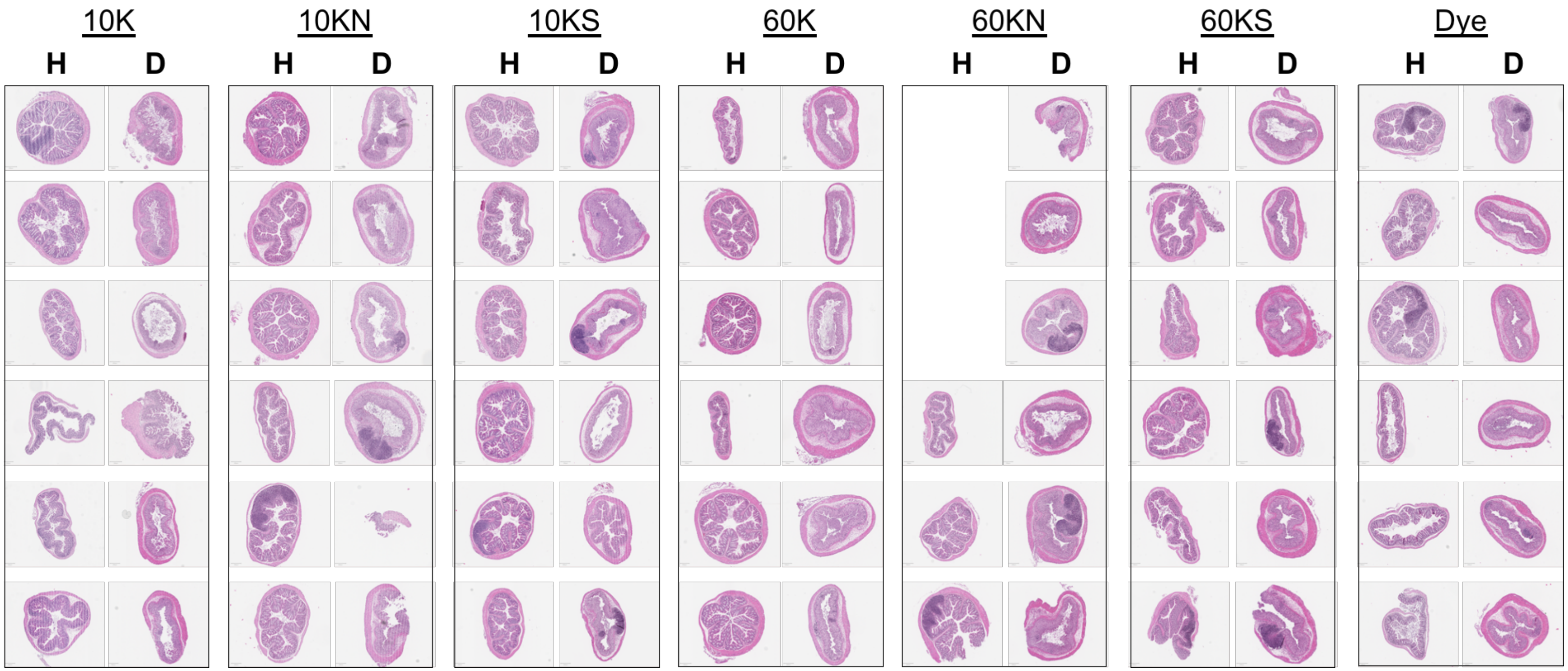


**Fig. S12. Representative histology images for individual mice in the Control and DSS-treated groups.** All mice were included for each polymer evaluation. H: Healthy mice; D: Mice with DSS-induced colitis. The histology images were scanned at 20x magnification (Aperio Slide Scanner, Leica Biosystems).

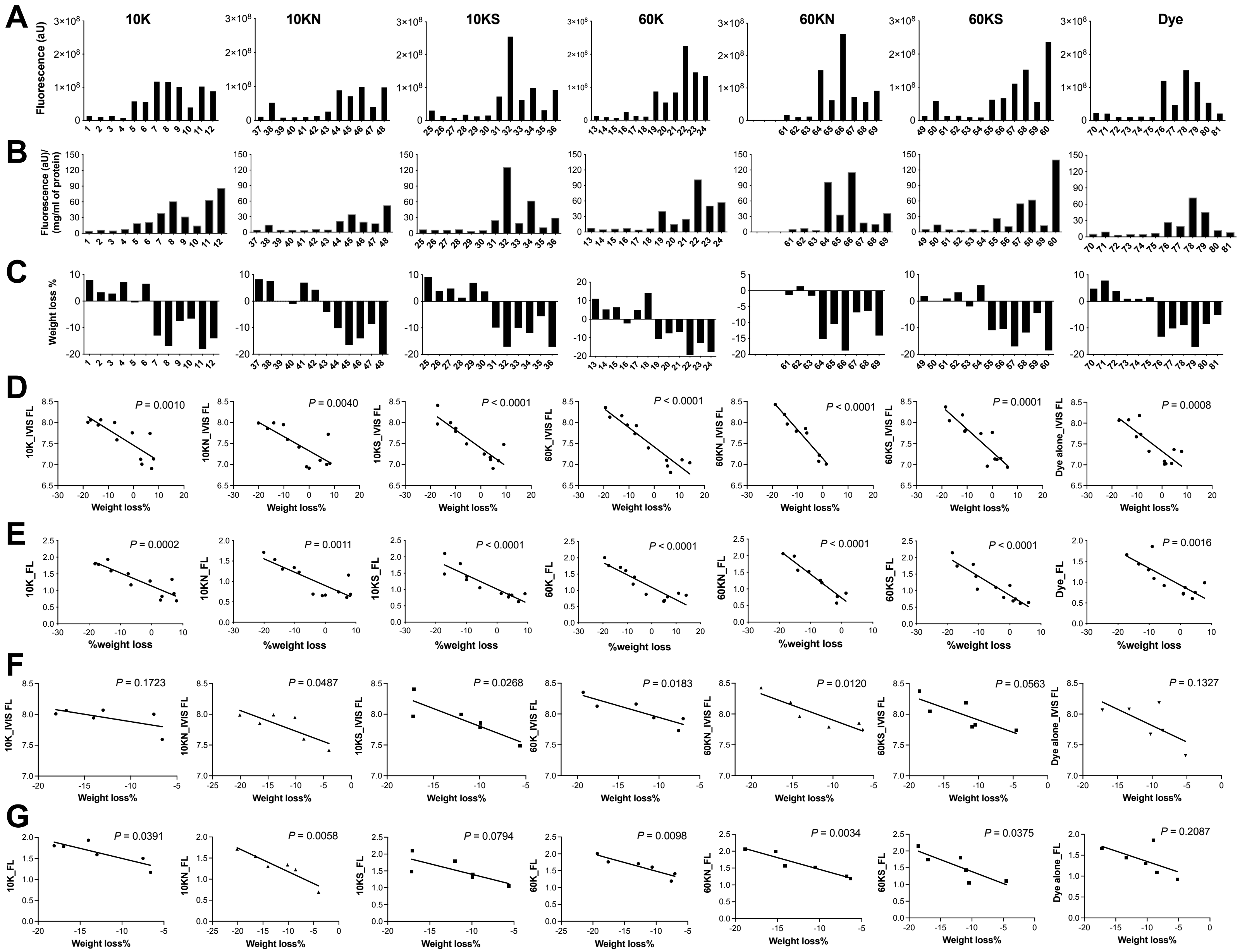
A
B
C
D
E
F
G
10K
10KN
10KS
60K
60KN
60KS
Dye
Fluorescence (aU)
Fluorescence (aU)/ (mg/ml of protein)
Weight loss %
Weight loss%
%weight loss
P = 0.0010
P = 0.0040
P < 0.0001
P < 0.0001
P < 0.0001
P = 0.0001
P = 0.0008
P = 0.0002
P = 0.0011
P < 0.0001
P < 0.0001
P < 0.0001
P < 0.0001
P = 0.0016
P = 0.1723
P = 0.0487
P = 0.0268
P = 0.0183
P = 0.0120
P = 0.0563
P = 0.1327
P = 0.0391
P = 0.0058
P = 0.0794
P = 0.0098
P = 0.0034
P = 0.0375
P = 0.2087

**Fig. S13. Correlations between fluorescence retention and weight loss% of individual mice in different treatment groups. (A)** Fluorescence was quantified by IVIS imaging of individual mice. **(B)** Fluorescence was quantified by homogenizing the colon tissue and normalized against the total protein in the colon tissue. **(C)** The body weight loss% of individual mice for different treatment groups. In (**A-C**), the x-axis indicates individual mice from #1 - #81. **(D)** The linear regression relationships between the log scale of the IVIS fluorescence (IVIS FL) and the body weight loss% of mice for all polymers and the dye alone. (**E**) The linear regression relationships between the log scale of the fluorescence in the homogenized colon tissue (FL) and the body weight loss% of mice for all polymers and the dye alone. (**F**) Correlations between the log scale of the IVIS fluorescence (IVIS FL) and the body weight loss% of colitic mice only for all polymers and the dye alone. (**G**) Correlations between the log scale of the fluorescence in the homogenized colon tissue (FL) and the body weight loss% of colitic mice only for all polymers and the dye alone.

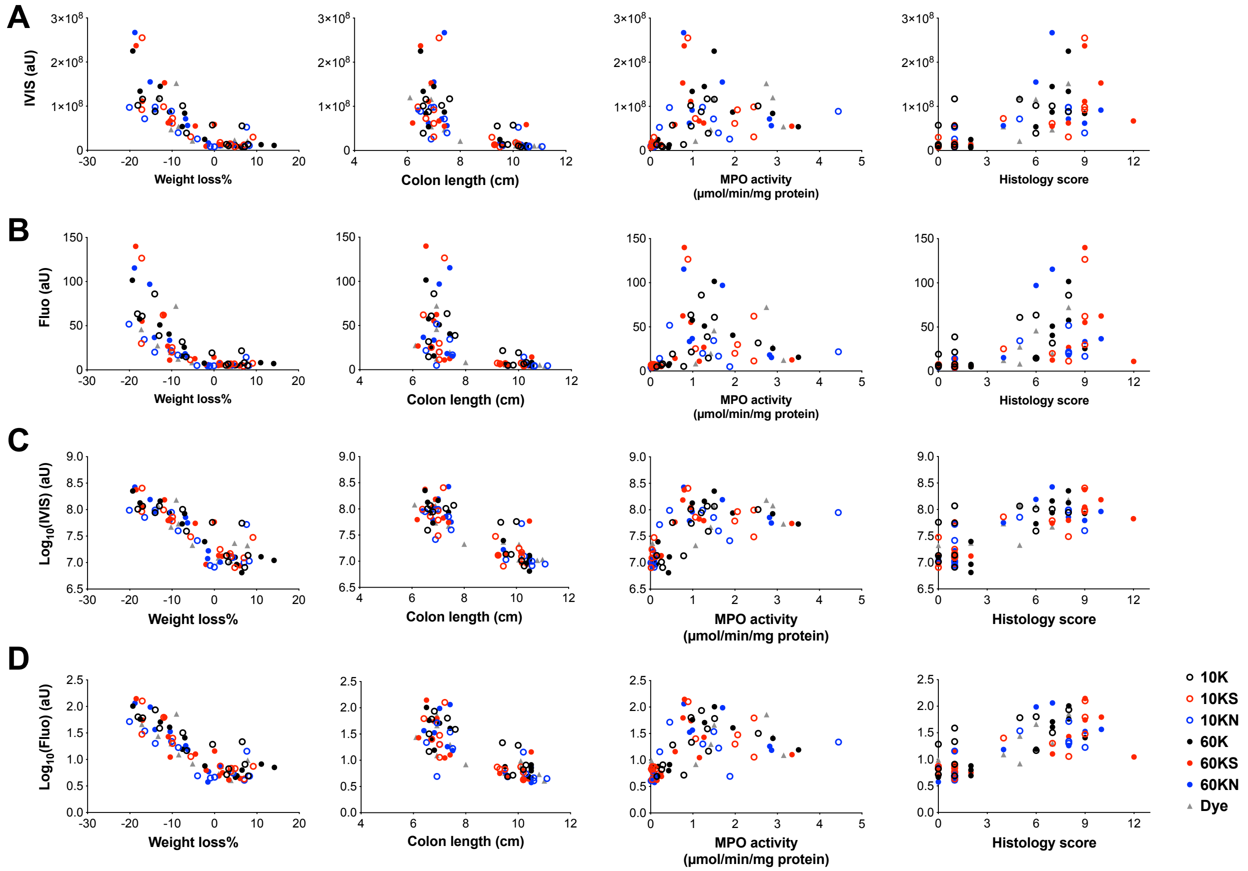


**Fig. S14. Correlations between fluorescence retention and weight loss%, colon length, MPO activity, and histology scores for all mice.** (**A**) Fluorescence quantified by IVIS imaging, denoted IVIS (aU), was plotted against body weight loss%, colon length, MPO activity, and histology scores for all polymers and the dye alone groups. (**B**) Fluorescence quantified by homogenizing colon tissue, denoted Fluo (aU), was plotted against body weight loss%, colon length, MPO activity, and histology scores for all polymers and the dye alone groups. (**C, D**) The log plots of fluorescence in (**A**) and (**B**), respectively. Data for all individual mice in different treatment groups were pooled together.

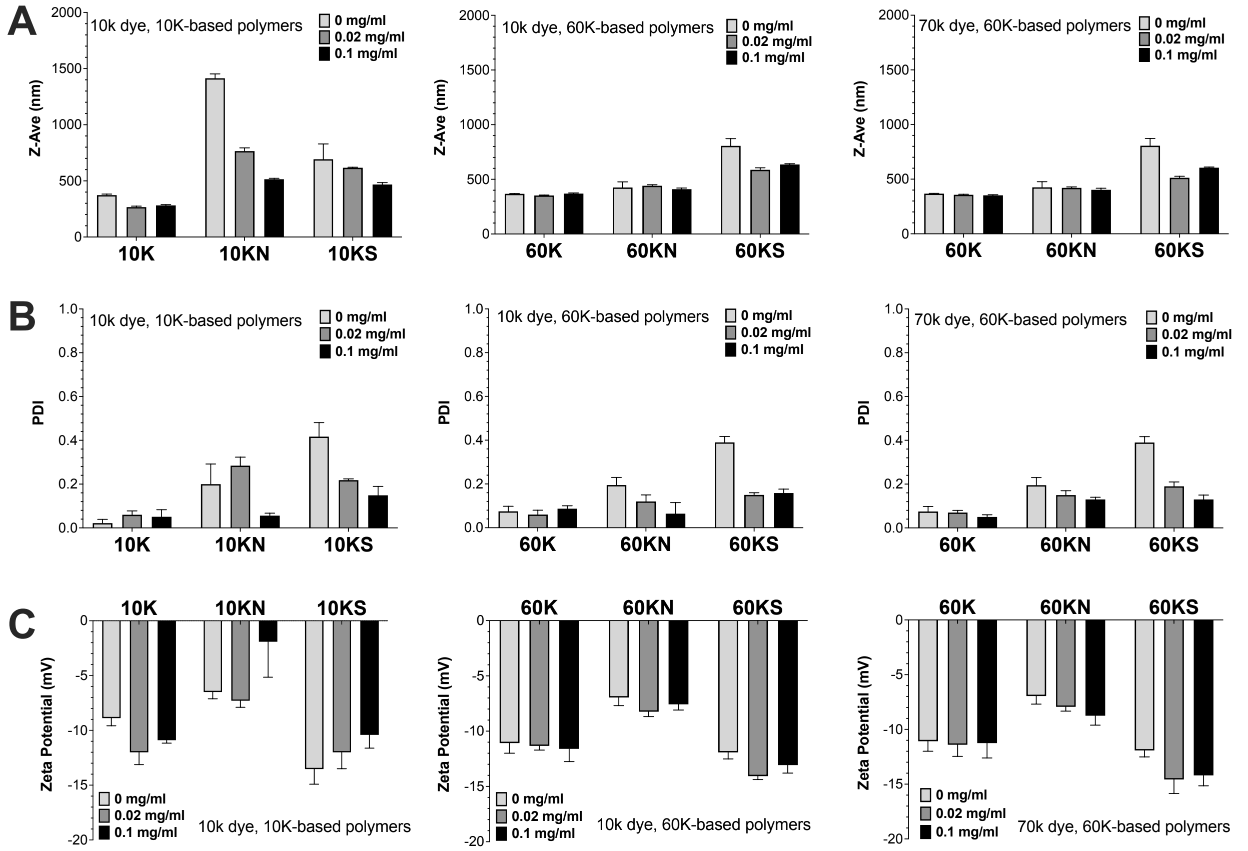


**Fig. S15. Different molecular weights and concentrations of the dye on the microgels' size, PDI, and zeta potential at 37°C.** (**A**) Size measurement of the polymers with different concentrations of the dye. Each polymer was at its concentration of $T_{cp}$ = 37°C. For 10K, 10KN, and 10KS, the concentration of each polymer was 5.6, 32.4, and 38.7 mg/ml. For 60K, 60KN, and 60KS, the concentration of each polymer was 4.9, 10.1, and 16.5 mg/ml. The dye concentration used was 0, 0.02, and 0.1 mg/ml. The molecular weight (MW) of the dye was indicated on each panel. (**B**) Comparison of the PDI for all polymers with the dye of different MWs and concentrations. **(C)** Comparison of the zeta potential for all polymers with the dye of different MWs and concentrations.

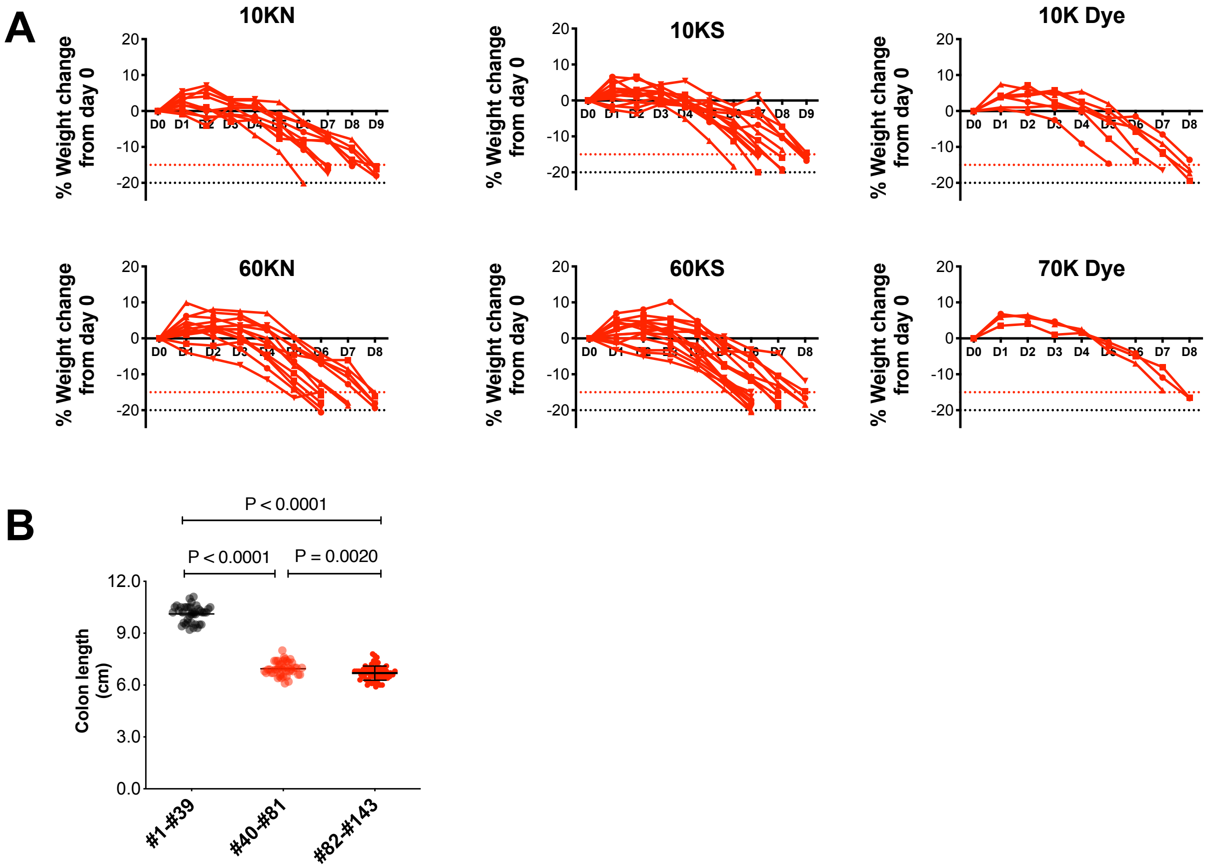


**Fig. S16. Weight loss% and colon length for *in vivo* evaluation of 10KN vs. 10KS using 10k dye as a control and 60KN vs. 60KS using 70k dye as a control. (A)** The body weight loss% of individual mice used for polymer evaluation *in vivo* in this study (mice #82 - #143). **(B)** Comparison of the colon length. #1 - #39 were healthy mice, and #40 - #81 were colitic mice; both were from Fig. S6 for comparison with mice #82 - #143 in this study. Note that both batches of mice, #40 - #81 and #82 - #143 were DSS-treated mice and they showed a significantly shorter colon length than the colon from #1 - #39 healthy mice ($P < 0.0001$). Further, mice #82 - #143 showed a shorter colon length than #40 - #81 ($P = 0.0020$), because mice #82 - #143 had a higher weight loss% than #40 - #81. Data are Mean ± SD, and Turkey's multiple comparison test is used for comparing multiple groups.

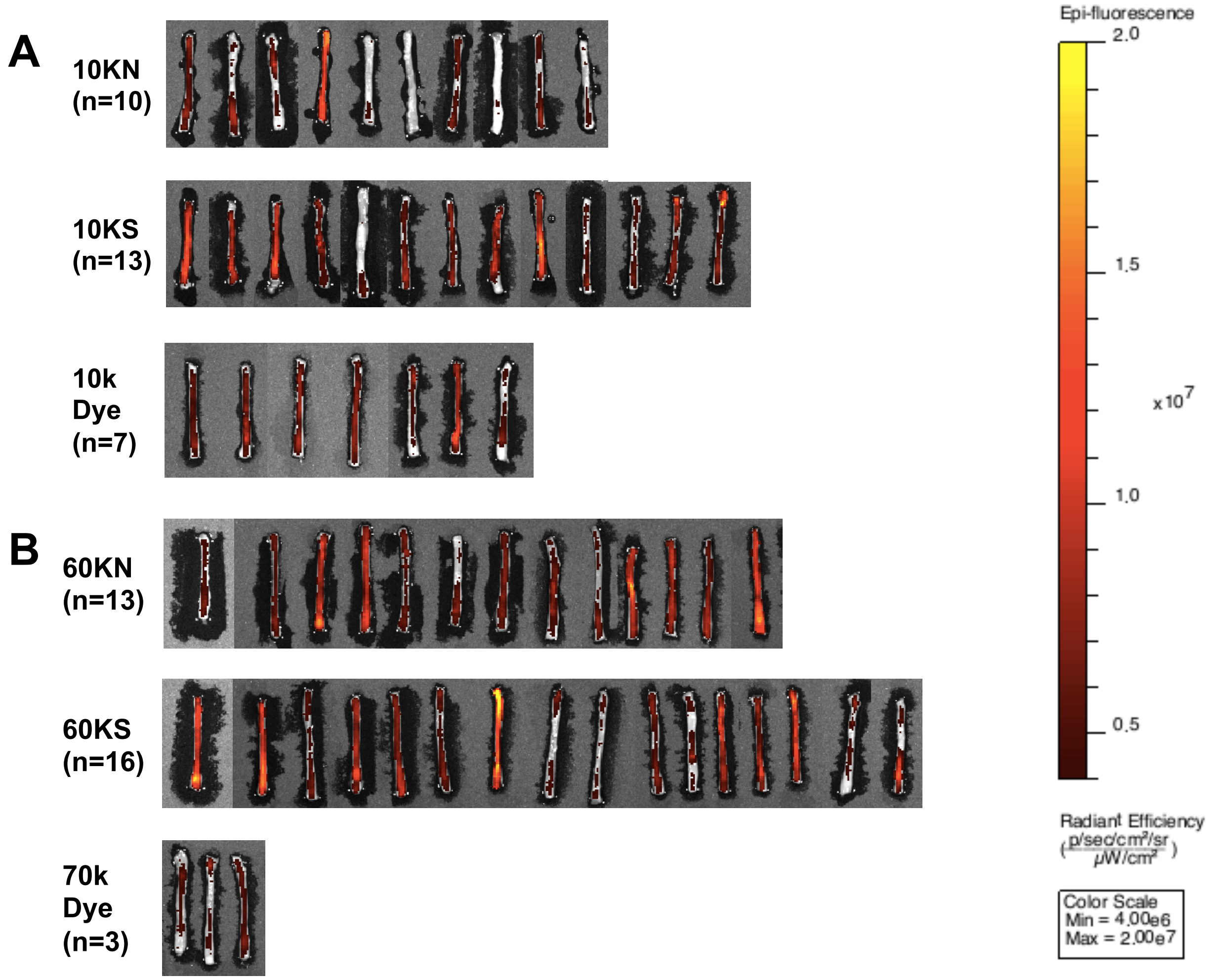


**Fig. S17. IVIS images for *in vivo* evaluation of 10KN vs. 10KS using 10k dye as a control and 60KN vs. 60KS using 70k dye as a control.** Since the IVIS imaging was performed when the body weight loss% of mice reached -15%, these studies were performed on different days on the same IVIS machine. The images were adjusted to the same scale of radiant efficiency for display.

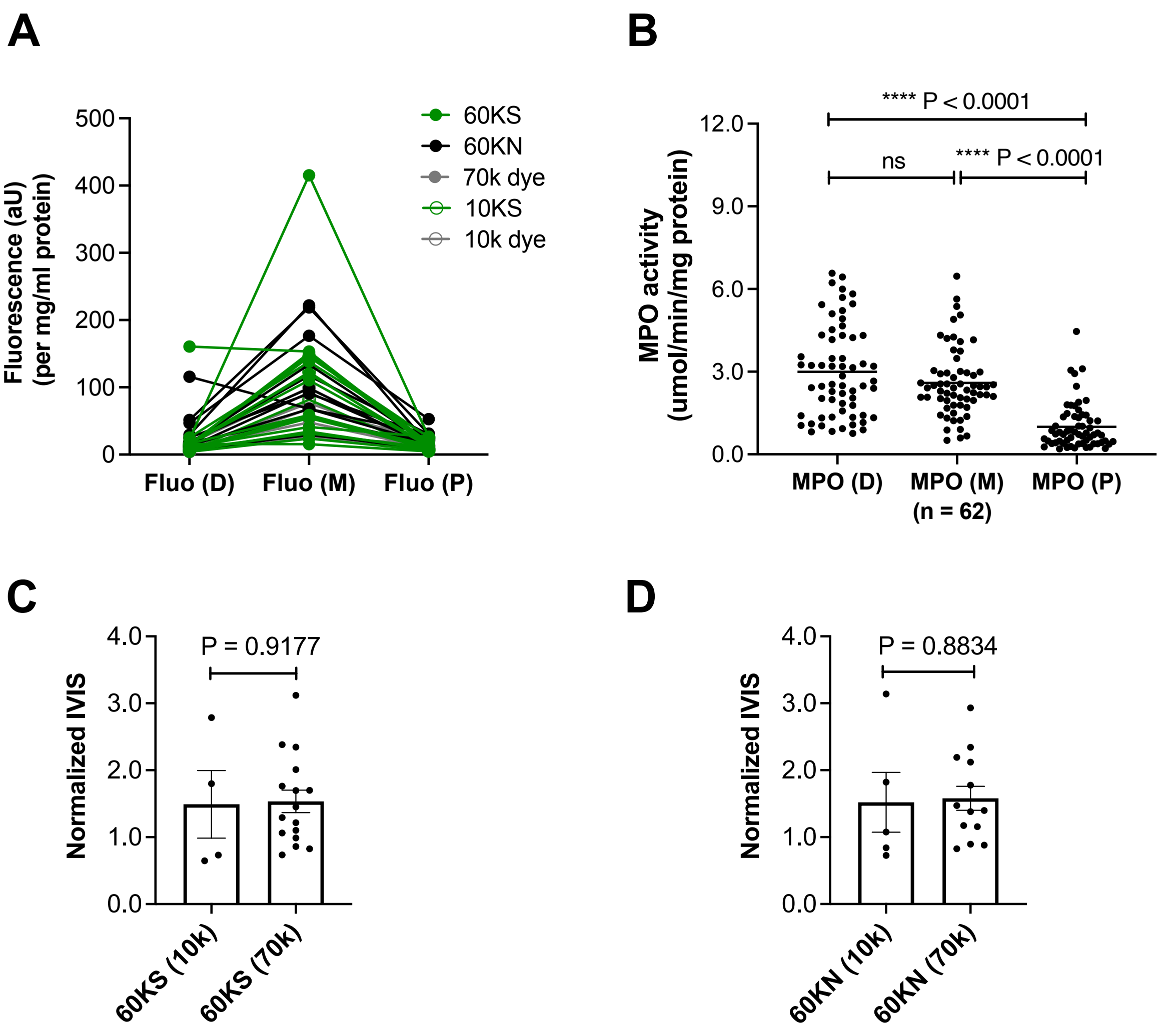


**Fig. S18. Molecular weight of the dye did not affect the fluorescence retention for 60KS and 60KN. (A, B)** Colon tissue heterogeneity in mice with colitis. Fluorescence retention in different colon sections (D: distal, M: middle, and P: proximal) was compared in (**A**). The MPO activity in different colon sections from individual mice was compared in (**B**). The distal and the middle colon showed a higher MPO activity than the proximal colon ($P < 0.0001$). **(C)** Comparison of the 10k and 70k dyes on the fluorescence retention of 60KS (n = 4 for 60KS with the 10k dye and n = 16 for 60KS with the 70k dye; $P = 0.9177$). **(D)** Comparison of the 10k and 70k dyes on the fluorescence retention of 60KN (n = 5 for 60KN with the 10k dye and n = 13 for 60KN with the 70k dye; $P = 0.8834$). Normalized IVIS was calculated by the IVIS fluorescence in the polymer groups divided by the IVIS fluorescence in the corresponding dye alone group.

| | Mouse # | IVIS | Normalized IVIS | MPO (P) | MPO (M) | MPO (D) | Fluo(P) | Fluo (M) | Fluo(D) |
|---|---|---|---|---|---|---|---|---|---|
| 10KN | 82 | 1.56e+007 | 0.49 | 0.71 | 1.79 | 3.55 | | 27.92 | |
| 10KN | 83 | 1.27e+007 | 0.39 | 0.86 | 2.17 | 3.48 | | 14.95 | |
| 10KN | 84 | 3.22e+007 | 1.00 | 1.96 | 1.32 | 0.76 | | 62.25 | |
| 10KN | 85 | 6.63e+007 | 2.06 | 1.48 | 5.37 | 1.85 | | 273.48 | |
| 10KN | 86 | 2.83e+007 | 0.88 | 0.39 | 2.59 | 5.10 | | 62.58 | |
| 10KN | 87 | 1.57e+007 | 0.49 | 0.56 | 2.14 | 6.57 | | 18.05 | |
| 10KN | 88 | 2.70e+007 | 0.84 | 0.43 | 2.24 | 2.85 | | 50.10 | |
| 10KN | 89 | 2.93e+007 | 0.91 | 0.25 | 2.41 | 6.23 | | 24.70 | |
| 10KN | 90 | 2.56e+007 | 0.80 | 0.21 | 1.38 | 4.24 | | 23.39 | |
| 10KN | 91 | 1.54e+007 | 0.48 | 0.37 | 1.70 | 3.48 | | 17.92 | |
| 10KS | 92 | 6.18e+007 | 1.92 | 1.33 | 5.64 | 4.32 | | 81.75 | |
| 10KS | 93 | 6.39e+007 | 1.99 | 0.40 | 3.04 | 2.49 | | 122.68 | |
| 10KS | 94 | 5.02e+007 | 1.56 | 1.89 | 2.56 | 1.16 | | 145.35 | |
| 10KS | 95 | 3.43e+007 | 1.07 | 0.77 | 2.10 | 2.40 | | 76.58 | |
| 10KS | 96 | 3.55e+007 | 1.10 | 0.39 | 0.91 | 4.16 | | 30.54 | |
| 10KS | 97 | 2.97e+007 | 0.92 | 0.27 | 2.46 | 3.20 | | 37.35 | |
| 10KS | 98 | 3.37e+007 | 1.05 | 0.28 | 4.90 | 4.66 | | 63.56 | |
| 10KS | 99 | 4.22e+007 | 1.31 | 0.25 | 2.87 | 5.46 | | 36.15 | |
| 10KS | 100 | 2.25e+007 | 0.70 | 0.51 | 1.50 | 6.00 | | 16.60 | |
| 10KS | 101 | 2.18e+007 | 0.68 | 0.77 | 2.56 | 2.66 | | 28.43 | |
| 10KS | 102 | 2.31e+007 | 0.72 | 0.66 | 5.06 | 1.11 | 5.49 | 33.15 | 6.04 |
| 10KS | 103 | 3.71e+007 | 1.15 | 0.23 | 2.50 | 4.92 | 8.07 | 47.87 | 13.39 |
| 10KS | 104 | 3.93e+007 | 1.22 | 0.20 | 1.42 | 1.42 | 25.30 | 74.54 | 11.74 |
| 10k dye | 105 | 2.75e+007 | 0.86 | 0.72 | 1.78 | 1.32 | | 67.97 | |
| 10k dye | 106 | 3.12e+007 | 0.97 | 1.02 | 2.07 | 0.89 | | 99.47 | |
| 10k dye | 107 | 3.60e+007 | 1.12 | 0.48 | 1.67 | 1.35 | | 76.70 | |
| 10k dye | 108 | 3.72e+007 | 1.16 | 1.03 | 2.92 | 1.05 | | 70.18 | |
| 10k dye | 109 | 2.55e+007 | 0.79 | 0.47 | 0.66 | 0.81 | 21.62 | 26.09 | 4.67 |
| 10k dye | 110 | 4.56e+007 | 1.42 | 1.61 | 3.75 | 5.44 | 7.95 | 94.94 | 16.42 |
| 10k dye | 111 | 2.22e+007 | 0.69 | 0.84 | 2.55 | 2.42 | 12.85 | 27.88 | 8.98 |

**Table S1. Tabulated data for all 10K-based polymers evaluated *in vivo* in Fig. 4.** The listed data include the IVIS quantifications, IVIS normalization, fluorescence in the homogenized colon tissue (Fluo; P: proximal, M: middle, and D: distal, indicating different sections of the colon tissue), and MPO data for 10KN, 10KS, and the 10k dye. All the Fluo and MPO data were normalized against the total protein in the homogenized colon tissue. Normalized IVIS was calculated by the IVIS fluorescence in the polymer groups divided by the IVIS fluorescence in the dye alone group.

| | Mouse # | IVIS | Normalized IVIS | MPO (P) | MPO (M) | MPO (D) | Fluo(P) | Fluo (M) | Fluo(D) |
|---|---|---|---|---|---|---|---|---|---|
| 60KN | 112 | 2.31e+007 | 0.90 | 0.82 | 2.50 | 1.33 | 16.66 | 68.21 | 115.88 |
| 60KN | 113 | 3.02e+007 | 1.17 | 3.11 | 3.01 | 1.34 | 28.12 | 90.45 | 25.27 |
| 60KN | 114 | 5.65e+007 | 2.19 | 1.37 | 0.88 | 3.25 | 52.79 | 176.76 | 46.75 |
| 60KN | 115 | 6.04e+007 | 2.34 | 2.93 | 3.79 | 3.22 | 14.46 | 221.94 | 29.29 |
| 60KN | 116 | 2.98e+007 | 1.16 | 1.03 | 2.02 | 2.54 | 12.17 | 98.68 | 19.36 |
| 60KN | 117 | 2.27e+007 | 0.88 | 1.21 | 2.45 | 1.58 | 10.18 | 29.06 | 7.40 |
| 60KN | 118 | 3.56e+007 | 1.38 | 1.44 | 4.26 | 4.33 | 14.51 | 92.04 | 11.05 |
| 60KN | 119 | 3.61e+007 | 1.40 | 2.47 | 1.95 | 1.07 | 7.22 | 30.96 | 4.41 |
| 60KN | 120 | 2.13e+007 | 0.83 | 0.74 | 2.30 | 5.82 | 31.26 | 67.77 | 13.53 |
| 60KN | 121 | 5.47e+007 | 2.12 | 0.27 | 2.96 | 2.03 | 30.30 | 117.09 | 16.75 |
| 60KN | 122 | 4.58e+007 | 1.78 | 0.49 | 2.08 | 1.78 | 16.15 | 134.07 | 16.68 |
| 60KN | 123 | 3.79e+007 | 1.47 | 0.48 | 2.06 | 5.22 | 16.16 | 55.08 | 17.72 |
| 60KN | 124 | 7.56e+007 | 2.93 | 0.96 | 2.16 | 2.29 | 30.69 | 218.90 | 51.41 |
| 60KS | 125 | 6.05e+007 | 2.35 | 0.51 | 1.96 | 4.53 | 14.18 | 153.44 | 160.78 |
| 60KS | 126 | 6.15e+007 | 2.38 | 1.22 | 4.33 | 2.30 | 26.35 | 138.05 | 25.28 |
| 60KS | 127 | 2.74e+007 | 1.06 | 1.78 | 2.79 | 0.83 | 10.94 | 54.39 | 7.37 |
| 60KS | 128 | 4.55e+007 | 1.76 | 1.80 | 2.34 | 1.03 | 17.59 | 151.00 | 14.59 |
| 60KS | 129 | 4.38e+007 | 1.70 | 1.36 | 4.16 | 3.23 | 12.14 | 120.23 | 16.58 |
| 60KS | 130 | 2.84e+007 | 1.10 | 0.70 | 2.99 | 3.20 | 8.16 | 58.09 | 8.89 |
| 60KS | 131 | 8.05e+007 | 3.12 | 1.43 | 2.22 | 1.40 | 26.56 | 415.43 | 22.65 |
| 60KS | 132 | 2.13e+007 | 0.83 | 4.46 | 6.46 | 3.28 | 7.58 | 34.38 | 5.56 |
| 60KS | 133 | 1.89e+007 | 0.73 | 3.05 | 2.00 | 2.04 | 14.28 | 54.50 | 8.51 |
| 60KS | 134 | 3.33e+007 | 1.29 | 0.80 | 2.41 | 2.47 | 26.41 | 41.18 | 11.38 |
| 60KS | 135 | 2.55e+007 | 0.99 | 0.62 | 4.09 | 0.93 | 6.32 | 23.93 | 4.29 |
| 60KS | 136 | 4.37e+007 | 1.70 | 1.76 | 4.21 | 1.99 | 12.43 | 110.56 | 16.76 |
| 60KS | 137 | 3.75e+007 | 1.45 | 0.88 | 2.53 | 5.69 | 11.77 | 58.81 | 16.35 |
| 60KS | 138 | 5.18e+007 | 2.01 | 0.38 | 2.96 | 3.15 | 11.36 | 149.11 | 14.87 |
| 60KS | 139 | 2.22e+007 | 0.86 | 0.72 | 1.24 | 4.32 | 10.51 | 31.65 | 8.36 |
| 60KS | 140 | 3.13e+007 | 1.21 | 0.42 | 0.51 | 6.44 | 4.91 | 15.64 | 15.96 |
| 70k dye | 141 | 2.52e+007 | 0.98 | 0.61 | 1.22 | 3.19 | 10.45 | 26.15 | 6.48 |
| 70k dye | 142 | 2.35e+007 | 0.91 | 0.39 | 0.60 | 2.49 | 16.67 | 24.02 | 6.13 |
| 70k dye | 143 | 2.86e+007 | 1.11 | 0.39 | 3.48 | 2.21 | 10.06 | 47.77 | 11.18 |

**Table S2. Tabulated data for all 60K-based polymers evaluated *in vivo* in Fig. 4.** The listed data include the IVIS quantifications, IVIS normalization, fluorescence in the homogenized colon tissue (Fluo; P: proximal, M: middle, and D: distal, indicating different sections of the colon tissue), and MPO data for 60KN, 60KS, and the 70k dye. All the Fluo and MPO data were normalized against the total protein in the homogenized colon tissue. Normalized IVIS was calculated by the IVIS fluorescence in the polymer groups divided by the IVIS fluorescence in the dye alone group.
